# Systems genetics uncover new loci containing functional gene candidates in *Mycobacterium tuberculosis*-infected Diversity Outbred mice

**DOI:** 10.1101/2023.12.21.572738

**Authors:** DM Gatti, AL Tyler, JM Mahoney, GA Churchill, B Yener, D Koyuncu, MN Gurcan, MKK Niazi, T Tavolara, AC Gower, D Dayao, E McGlone, ML Ginese, A Specht, A Alsharaydeh, PA Tessier, SL Kurtz, K Elkins, I Kramnik, G Beamer

## Abstract

*Mycobacterium tuberculosis,* the bacillus that causes tuberculosis (TB), infects 2 billion people across the globe, and results in 8-9 million new TB cases and 1-1.5 million deaths each year. Most patients have no known genetic basis that predisposes them to disease. We investigated the complex genetic basis of pulmonary TB by modelling human genetic diversity with the Diversity Outbred mouse population. When infected with *M. tuberculosis*, one-third develop early onset, rapidly progressive, necrotizing granulomas and succumb within 60 days. The remaining develop non-necrotizing granulomas and survive longer than 60 days. Genetic mapping using clinical indicators of disease, granuloma histopathological features, and immune response traits identified five new loci on mouse chromosomes 1, 2, 4, 16 and three previously identified loci on chromosomes 3 and 17. Quantitative trait loci (QTLs) on chromosomes 1, 16, and 17, associated with multiple correlated traits and had similar patterns of allele effects, suggesting these QTLs contain important genetic regulators of responses to *M. tuberculosis*. To narrow the list of candidate genes in QTLs, we used a machine learning strategy that integrated gene expression signatures from lungs of *M. tuberculosis-*infected Diversity Outbred mice with gene interaction networks, generating functional scores. The scores were then used to rank candidates for each mapped trait in each locus, resulting in 11 candidates: *Ncf2, Fam20b, S100a8, S100a9, Itgb5, Fstl1, Zbtb20, Ddr1, Ier3, Vegfa,* and *Zfp318*. Importantly, all 11 candidates have roles in infection, inflammation, cell migration, extracellular matrix remodeling, or intracellular signaling. Further, all candidates contain single nucleotide polymorphisms (SNPs), and some but not all SNPs were predicted to have deleterious consequences on protein functions. Multiple methods were used for validation including (i) a statistical method that showed Diversity Outbred mice carrying PWH/PhJ alleles on chromosome 17 QTL have shorter survival; (ii) quantification of S100A8 protein levels, confirming predicted allele effects; and (iii) infection of C57BL/6 mice deficient for the *S100a8* gene. Overall, this work demonstrates that systems genetics using Diversity Outbred mice can identify new (and known) QTLs and new functionally relevant gene candidates that may be major regulators of granuloma necrosis and acute inflammation in pulmonary TB.

## INTRODUCTION

The number of humans who develop active pulmonary tuberculosis (TB) is small compared to those who eliminate or control *Mycobacterium tuberculosis* (5-10% *vs* 90-95%), yet morbidity and mortality from TB remain high. Although COVID-19 mortality temporarily surpassed global TB mortality, TB has remained in the top two leading causes of death due to an infectious disease for decades, killing more people than HIV/AIDS and malaria. Pulmonary TB is the most common and most contagious form of TB, with mortality rates >40% if untreated or if caused by antibiotic resistant *M. tuberculosis* [1-8]. Human responses to *M. tuberculosis* infection range from fulminant pulmonary TB that develops within weeks to lifelong control of latent infection or complete clearance of bacilli [9-11]. Further, a body of evidence shows an interesting paradox: Immune competence is necessary to restrict *M. tuberculosis* growth [12], but is not sufficient to prevent disease [13].

The variable responses to *M. tuberculosis* and lack of single genetic defects in most patients indicate a complex genetic basis for pulmonary TB, and this has been investigated by linkage association mapping, genome-wide association studies, and other methods, recently reviewed [14-17]. These reviews frequently identify knowledge gaps attributable to the observations that the most used laboratory mouse strains do not replicate key disease traits (e.g., granuloma necrosis) of human pulmonary TB [18-23]. To address these gaps, we and others use the Diversity Outbred mouse population and Collaborative Cross recombinant inbred strains [24-26], some of which do develop human-like pulmonary TB following *M. tuberculosis* infection. These mice provide valuable resources to model complex genotype-phenotype associations; tools to dissect the genetic basis of disease; and a means to test for effects of candidate genetic polymorphisms *in vivo*.

The Diversity Outbred mouse population originated by breeding eight inbred founder strains together, resulting in an experimental population with balanced allele frequencies of one-eighth across the genome [27]. This is important for genetic mapping studies because low allele frequencies in natural populations can diminish power and increase false positive findings [28]. Further, Diversity Outbred mice carry over 40 million variants [29], some of which alter regulatory elements, splice sites, and protein-coding sequences. This defined genetic architecture allows rigorous investigation of genotype-phenotype association in context of *M. tuberculosis* infection.

Here, to find genetic loci associated with pulmonary TB, we used quantitative trait locus (QTL) mapping. Next we ranked candidates genes within the *Diversity Outbred tuberculosis susceptibility* (*Dots*) loci that were associated with correlated, colocalized traits by using a machine learning algorithm [30, 31] to find genes functionally related to the mapped traits and the fit models scored each candidate [32]. All candidates contain a variety of SNPs as annotated in Mouse Variation Registry (MVAR). Seven of the eleven candidates contain missense SNPs in protein coding regions, and of those, the SNPs in four candidates (*S100a8, Itgb5, Fstl1, and Zfp318)* are predicted to have deleterious consequences on protein functions.

Published studies have shown three candidates (*Itgb5, Fstl1, S100a8*) involved in bacterial lung diseases that includes *in vitro*, or *in vivo M. tuberculosis* infection [33-39]. The other eight candidates have no known roles in *M. tuberculosis* infection but have been shown in other systems to contribute to cell stress responses, signaling pathways, adhesion and migration; extracellular matrix synthesis, tissue remodeling and angiogenesis; immune cell metabolism; macrophage inflammatory responses; and viral hepatitis [40-52]. Overall, ten candidate genes have roles in innate immune responses suggesting that genetically controlled responses of epithelial and endothelial cells, neutrophils, and monocytes, macrophages to *M. tuberculosis* bacilli are the primary drivers of susceptibility to *M. tuberculosis* and to disease progression in pulmonary TB. Only 1 candidate has a direct role in acquired, antigen-specific immunity.

## METHODS

### Ethics Statement

Tufts University’s Institutional Animal Care and Use Committee (IACUC) approved this work under protocols G2012-53; G2015-33; G2018-33; and G2020-121. Tufts University’s Institutional Biosafety Committee (IBC) approved this work under registrations: GRIA04; GRIA10; GRIA17, and 2020-G61.

### Mice

Female Diversity Outbred mice (n=850) from generations 15 16, 21, 22, 34, 35, 37 and 42 and the inbred founder strains: A/J, C57BL/6J, 129S1/SvlmJ), NOD/LtJ, NZO/HILtJ, CAST/EiJ, PWK/PhJ, and WSB/EiJ mice (n=15-59 per strain) were purchased from The Jackson Laboratory (Bar Harbor, ME) and group housed (n=5-7 mice per cage) on Innovive (San Diego, CA) or Allentown Inc (Allentown, NJ) ventilated, HEPA-filtered racks in the New England Regional Biosafety Laboratory (Tufts University, Cummings School of Veterinary Medicine, North Grafton, MA) or at The Ohio State University Columbus, OH. The light cycle was 12 hours of light; 12 hours of dark. Two breeding pairs of female and male C57BL/6 inbred mice carrying null mutation for *S100a8* gene were a kind gift of Dr. Philippe Tessier, Department of Microbiology and Immunology, Faculty of Medicine, Université Laval. After quarantine, breeders were used to establish a colony of S100a8 homozygous knock out (KO), heterozygous (HET) and wild-type (WT) C57BL/6 inbred mice. Mice were housed in disposable sterile caging or re-usable autoclaved caging containing sterile corn-cob bedding, with sterile paper nestlets (Scotts Pharma Solutions, Marlborough, MA), and/or sterile enrichment paperboard or plastic “houses”. Cages were changed every other week or sooner if soiled. Mice were provided with sterile mouse chow (Envigo, Indianapolis, IA) and sterile, acidified water *ad libidum*.

### *M. tuberculosis* Aerosol Infection

Female Diversity Outbred mice and inbred founder strains were infected with aerosolized *M. tuberculosis* strain Erdman bacilli using a custom-built CH Technologies system [24, 39, 53] or a Glas-col (Terre Haute, ID) system [54, 55] between eight and ten weeks of age. Male and female C57BL/6 *S100a8* KO, HET, and WT mice were infected between eight and sixteen weeks of age. For each aerosol infection, the retained lung dose was determined by euthanizing a cohort of four to twelve mice 24 hours after exposure, homogenizing the entire lungs in 5mL sterile phosphate buffered saline, and plating the entire homogenate onto OADC-supplemented 7H11 agar. After 3-4 weeks at 37°C, *M. tuberculosis* colony forming units were counted. Mice were infected with ∼100 colony forming units in the first two experiments, and ∼25 colony forming units in the subsequent eight experiments.

### Quantification of TB-related Traits (Phenotyping)

#### Survival

IACUC protocols disallowed natural death as an endpoint. Therefore, as a proxy of survival, we used the day of euthanasia due to any single criterion: Severe weakness/lethargy; or respiratory distress; or body condition score < 2 [56]. We confirmed morbidity was due to pulmonary TB by finding: (i) Large nodular, or severe diffuse lung lesions; (ii) histopathology confirmation of severe granulomatous lung infiltrates; (iii) growth of viable *M. tuberculosis* colonies from lung tissue; and (iv) absence of other diseases. Twenty-one *M. tuberculosis* infected Diversity Outbred mice were excluded due to co-morbidity that developed during the experiment.

#### Weight loss

Mice were weighed 1 to 3 days prior to *M. tuberculosis* aerosol infection, at least once per week during infection, and immediately before euthanasia. For each mouse, weight loss was calculated as the percent loss from peak body weight.

#### Lung granuloma necrosis

Immediately after euthanasia, lung lobes were removed and inflated and fixed in 10% neutral buffered formalin (5-10 mL per lobe), processed, and embedded in paraffin, sectioned at 5µm, and stained with hematoxylin and eosin with or without carbol fuschin for acid-fast bacilli at Tufts University, Cummings School of Veterinary Medicine, Core Histology Laboratory (North Grafton, MA). Hematoxylin and eosin-stained glass slides were magnified 400 times and digitally scanned by Aperio, LLC (Sausalito, CA) ScanScope scanners at 0.23 microns per pixel at The Ohio State University’s Comparative Pathology and Mouse Phenotyping Shared Resources Core resource (Columbus, OH) or by Aperio, LLC (Sausalito, CA) AT2 scanners at 0.23 microns per pixel at Vanderbilt University Medical Center’s Digital Histology Shared Resource (Nashville, TN). Lung granuloma necrosis was quantified in one lung lobe per mouse by our previously validated, deep learning image analysis method [57] and reported here as a ratio of granuloma necrosis per lung tissue area.

#### M. tuberculosis lung burden

Immediately after euthanasia, 2 or 3 lung lobes were removed from each mouse and homogenized in sterile phosphate buffered saline (1mL per lobe), serially diluted, plated onto OADC-supplemented 7H11 agar, incubated at 37°C for 3-4 weeks, after which colonies were counted, and *M. tuberculosis* lung burden in the lungs was calculated as described [58].

#### Lung cytokines and chemokines

Lung homogenates were stored at ™80°C until the experiment ended. Lung homogenates were then thawed overnight at 4° serially diluted and tested for CXCL5, CXCL2, CXCL1, tumor necrosis factor (TNF), matrix metalloproteinase 8 (MMP8), S100A8, interferon-gamma (IFN-γ), interleukin (IL)-12p40, I-L12p70, IL-10, and vascular endothelial growth factor (VEGF) by sandwich ELISA using antibody pairs and standards from R&D Systems (Minneapolis, MN), Invitrogen (Carlsbad, CA), eBioscience (San Diego, CA), or BD Biosciences (San Jose, CA, USA), per kit instructions. Lung homogenate ELISA results from five of the experiments using Diversity Outbred mice have been published and analyzed for biomarkers previously [39].

### Phenotype Correlation

We took the log of each phenotype after adding one (to ensure that zero was not converted to negative infinity) and regressed out the effect of the experimental batch. We then standardized the residuals and estimated the Pearson correlation between all pairs of phenotypes.

### Gene Expression

One lung lobe from 98 Diversity Outbred mice was homogenized in TRIzol, stored at ™80°C, and RNA was extracted using Pure Link mini-kits (Life Technologies, Carlsbad, CA). Boston University’s Microarray and Sequencing Resource Core Facility (Boston, MA) confirmed quality and quantity were sufficient for microarray analyses. Mouse Gene 2.0 ST CEL files were normalized to produce gene-level expression values using the implementation of the Robust Multiarray Average (RMA) in the Affy package (version 1.62.0) included in the Bioconductor software suite and an Entrez Gene-specific probeset mapping (17.0.0) from the Molecular and Behavioral Neuroscience Institute (Brainarray) at the University of Michigan. Array quality was assessed by computing Relative Log Expression (RLE) and Normalized Unscaled Standard Error (NUSE) using the affyPLM package (version 1.59.0). The CEL files were also normalized using Expression Console (build 1.4.1.46) and the default probesets defined by Affymetrix to assess array quality using an AUC metric computed from sets of negative and positive control probes; all samples used in this analysis had an AUC > 0.8. Moderated *t*-tests and ANOVAs were performed using the limma package (version 3.39.19) (i.e., creating simple linear models with lmFit, followed by empirical Bayesian adjustment with eBayes). Correction for multiple hypothesis testing was accomplished using the Benjamini-Hochberg false discovery rate (FDR). To remove microarray probes that intersected with Diversity Outbred SNPs, we intersected the Diversity Outbred founder strain SNPs [59] with the vendor-provided probes and removed probes containing SNPs. All microarray analyses were performed using the R environment for statistical computing (version 3.6.0). A related microarray dataset and other secondary analyses have been published elsewhere [39, 60] and deposited in Gene Expression Omnibus (GEO), and assigned Series ID GSE179417.

### Genotyping

We collected tail tips from each Diversity Outbred mouse and sent them to Neogen (Lincoln, NE) for DNA isolation and genotyping. Neogen genotyped the mice on the Illumina GigaMUGA platform, which contains 143,259 markers [61]. Genotypes of *S100a8* KO, HET, and WT C57BL/6 inbred mice were confirmed by polymerase chain reaction (TransnetYX, Cordova, TN).

### Haplotype Reconstruction and SNP Imputation

We used 137,302 GigaMUGA marker positions located on the autosomes and chromosome X found at https://github.com/kbroman/MUGAarrays/blob/main/UWisc/gm_uwisc_v1.csv and the R package *qtl2* to reconstruct the Diversity Outbred haplotypes using the founder and Diversity Outbred allele calls, and used the haplotype reconstructions to impute the founder SNPs onto the Diversity Outbred genomes [62].

### Quantitative Trait Locus Mapping

We included Diversity Outbred mice that survived *M. tuberculosis* infection for 250 days or less because age-related comorbidities began to appear and complicated interpretation. We used *qtl2* [62] to perform linkage mapping using the founder haplotypes and association mapping using the imputed SNPs. We calculated the kinship between mice using the leave-one-chromosome-out method, which excludes the current chromosome in kinship calculations [63]. We standardized each phenotype and mapped with the Diversity Outbred outbreeding generation as an additive covariate and used the linear mixed-effects model with one kinship matrix per chromosome. We estimated the genome-wide significance thresholds by permuting the samples 1,000 times and performed a genome scan with each permutation. We retained the maximum log_10_ of the odds ratio (LOD) score from each permutation and estimated the genome-wide significance threshold of 7.6 from the 95th percentile of the empirical distribution of maximum LOD scores under permutation. We estimated the support interval around each peak using the 95% Bayesian Credible Interval.

For each peak with a LOD score above the genome-wide threshold > 7.6, we then searched for peaks associated with other traits that had LOD scores > 6 and confidence intervals that overlapped [64]. Our rationale was that the probability that a peak is biologically relevant, given that another trait has a co-located peak, is higher than the probability that a peak is significant with no prior evidence.

### Candidate Gene Selection

Within each QTL interval, we imputed the founder SNPs onto the Diversity Outbred mouse genomes using *qtl2* and performed association mapping. We selected the SNPs that were within a 1 LOD drop of the peak SNP in the QTL interval and filtered them to retain ones with missense, splice, or stop codon effects as annotated by the Sanger Mouse Genome Project [59]. We considered the genes in which these polymorphisms occurred as candidate causal genes for the associated trait(s).

### Trait-related Gene Sets

Because causal variants within a QTL may exert their influence through mechanisms other than gene expression, identifying differentially expressed genes within the QTL may be insufficient for ranking causal genes. Here we took an alternative approach and ranked candidates in each QTL based on their predicted association with the mapped traits. To do this, we trained an SVM to classify trait-related genes, and then used the trained SVM to score each positional candidate gene as trait-related or not-trait-related. We defined the training set of trait-related genes for the SVM as those genes that were highly correlated to the measured trait using the gene expression data described above. We calculated the Pearson correlation between the abundance of each transcript, and each physiological trait using rank Z normalized gene expression and traits. For each trait, we defined the training set of trait-related genes as the 500 genes with the largest magnitude Pearson correlation to the trait. We have made these gene lists available as a set of zipped text files in Supplemental File 1.

### Support Vector Machine classifier training

We trained SVMs to classify genes in each gene list as trait-related using features derived from the Functional Network of Tissues in Mouse [32]. The nodes in this network are genes, and the edges between them are weights between 0 and 1 that predict the likelihood that each pair of genes is annotated to the same Gene Ontology (GO) term or KEGG pathway [30]. Values closer to one indicate more certainty that the genes are more likely to be annotated to the same GO term or KEGG pathway and thus functionally related. The weights were derived using Bayesian integration of data sets from numerous sources of data, including gene expression, protein-protein interaction data, and phenotype annotations [32]. We used the top edges of the mouse lung network downloaded on March 31, 2021, from http://fntm.princeton.edu.

### Application of Support Vector Machine classifiers to identify genes functionally related to traits

We used SVMs to classify each positional candidate as trait-related or not-trait-related, as described previously [30, 31]. Briefly, the expression-derived gene sets for each lung trait served as the *positive labeled set* of genes. We used the R package e1071 [65] to train SVMs to distinguish this set of genes from a balanced set of genes drawn randomly from the remaining genes in the lung network. The randomly selected genes were the *unlabeled set*. We performed this training 100 times, each time with a new set of random unlabeled genes. The SVMs were trained to distinguish positive labeled genes from unlabeled genes using the connection weights to the positive labeled genes. It is expected that positively labeled genes have relatively strong connections to each other because they are functionally related. It is further expected that randomly drawn genes will be unrelated to the trait and to the positive labeled set and will thus have relatively lower connection weights to the positive labeled genes. The SVM learns to distinguish these two groups of genes, and the resulting model can be used to classify genes that have not been seen before based on their connection weights to the positively labeled genes. We initialized each run by tuning the SVM over a series of cost parameters, starting with the sequence 1025 to 102 by factors of 10, and iteratively narrowing the range until we found a series of eight cost parameters that maximized accuracy. In running each SVM, we used a linear kernel and 10-fold cross-validation.

We calculated the area under the receiver operating characteristic curves (AUC) for each set of trait-related genes as follows. We defined labeled positives (LP) as positive labeled genes that were classified by the SVM as trait related. Unlabeled negatives (UN) were unlabeled genes that were classified by the SVM as not trait related. Unlabeled positives (UP) were unlabeled genes that were classified as trait-related, and labeled negatives (LN) were positive labeled genes that were classified as not trait-related. These terms are conceptually like true/false positive and true/false negative scores. However, because unlabeled genes may not be truly unrelated to the trait, we cannot call them true negatives. Instead, we call them *unlabeled*. We generated ROC curves using the *Unlabeled Predicted Positive Rate* (UPPR = UP/(UP+UN)), which is akin to the false positive rate, and the labeled positive rate (LPR = LP/(LP+UN)), which is akin to the false negative rate, along a series of SVM scores from the minimum to the maximum. We then calculated the average AUC across all 100 SVMs.

### Positional Candidate Scoring

After training SVMs for each trait, we scored all positional candidate genes in each QTL, defined as the minimum to the maximum position across a set of overlapping QTLs. Each candidate gene received one score for each trait that mapped to that location. To compare scores across traits, we used the UPPR for each gene at its calculated SVM score. The UPPR varies between 0 and 1, allowing us to compare scores for candidate genes across models. To visually compare across models, we used the −log10(UPPR) such that genes with very small UPPR (very high confidence) got large positive values. In contrast, the SVM scores cannot be used to compare across models because they are unbounded and vary from model to model. Within each pleiotropic QTL, each gene received a score from each trait that mapped to the QTL.

### Mouse Genome Build and Database Versions

We used mouse genome build GRCm38 and SNPs and Indels from the Sanger Mouse Genomes Project, version 7, which uses Ensembl version 97 gene models. We also used and cross-referenced candidates with the Mouse Phenome Database GenomeMUSter, the Mouse Genome Informatics databases [66], and Ensembl’s Variant Effect Predictor tool.

## RESULTS

### Survival and body weight changes

We infected Diversity Outbred mice by aerosol with ∼100 *M. tuberculosis* colony forming units in the first two experiments (N=167), and ∼25 colony forming units in the subsequent eight experiments (N=683). Infection reduced survival of Diversity Outbred mice compared to identically housed, age-, gender-, and generation-matched uninfected Diversity Outbred mice and to identically housed age- and gender-matched infected C57BL/6J inbred mice (Figure 1A). Approximately one-third of infected Diversity Outbred mice succumbed prior to 60 days (Figure 1A) reflecting early mortality between 20-56 days that peaked at 30 days (Figure 1B). This supersusceptible fraction of the Diversity Outbred population has been named Progressors [24, 57, 60, 67]. Morbidity in all Progressors was due to pulmonary TB, confirmed by histology, recovery of viable *M. tuberculosis* bacilli from the lungs and absence of other diseases. After the first mortality wave subsided, cumulative survival declined slowly to nearly 600 days with no discernable mortality waves (Figure 1A and 1B). This relatively resistant fraction of Diversity Outbred mice has been named Controllers [24, 57, 60, 67]. The eight founder strains survived at least 40 days of *M. tuberculosis* infection, without early mortality (Figure 1A).

**Figure 1.**
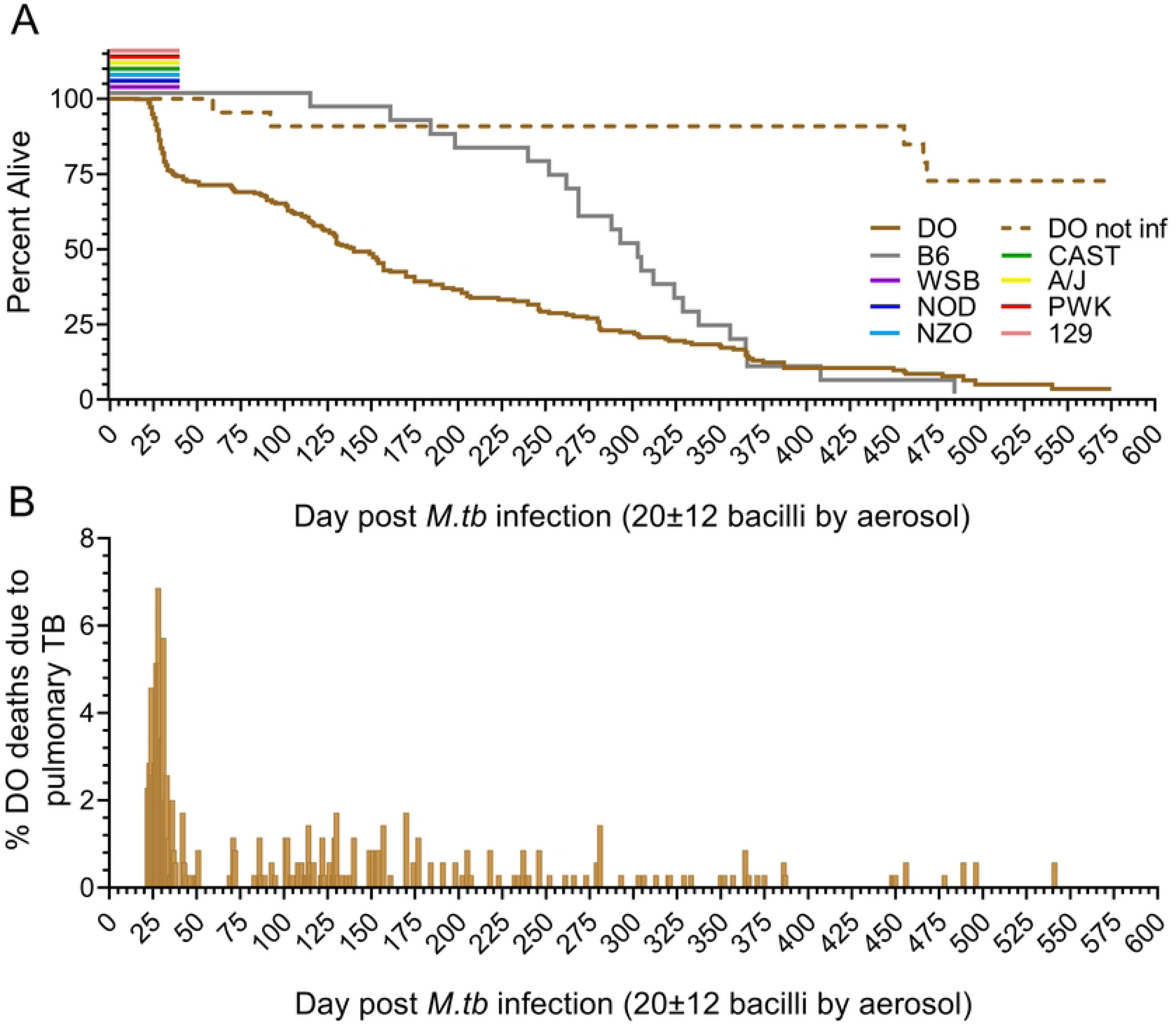
Mouse survival following exposure to a low dose of aerosolized *M. tuberculosis* strain Erdman. Diversity Outbred (DO) mice (n = 680, brown solid line), and the eight inbred founder strains (n = 15 to 78, colored lines) were infected with *M. tuberculosis* strain Erdman bacilli by aerosol. Panel A shows cumulative survival extending to nearly 600 days post infection. Approximately 30% of the DO population succumbed to pulmonary TB by 60 days post infection. Of the eight inbred founder strains, survival studies were completed for the C57BL/6J inbred strain; the other seven inbred founder strains were euthanized 40 days after *M. tuberculosis* infection. No inbred founder strain or non-infected (NI) DO mice (n = 53, dashed line) showed mortality within the same period. Panel B shows the daily mortality for *M. tuberculosis*-infected DO mice, highlighting the early wave of mortality that peaked between 25- and 35-days post infection.

All mice were weighed prior to infection, during infection, and immediately before euthanasia. Non-infected Diversity Outbred mice gained weight until they developed other diseases or were euthanized at the experiment end (Supplemental Figure 1A). Progressors gained weight for 2-3 weeks, and then quickly lost weight (Supplemental Figures 1B). Controllers and C57BL/6J inbred mice gained weight for long and variable durations through about 250 days of infection, and then most but not all slowly lost weight (Supplemental Figures 1C, 1D). We questioned whether pre-infection body weight influenced differential susceptibility. Retrospective analysis identified no significant differences in pre-infection body weights of non-infected Diversity Outbred mice compared to Progressors; and a significant difference (average of 1.75 gm lower) in mean body weights of non-infected Diversity Outbred mice and Progressors compared to Controllers (Supplemental Figure 2A). Whether this is spurious or biologically relevant (i.e., heavier pre-infection body weight partially protects) remains to be determined. Supplemental Figures 2B, 2C, and 2D show correlations between survival and eight clinical indicators of disease. Seven indicators positively correlate with survival, including pre-infection body weight which was weakly positive. Only one indicator, the rate of body weight loss, had negative correlations with survival and the duration of weight loss.

### Lung Histology and Automated Image Analysis of Granuloma Necrosis

By eight weeks of *M. tuberculosis* infection, Diversity Outbred mice showed a spectrum of lung lesions visible at low magnification (Figure 2) with variation in severity (minimal to marked); distribution of cellular infiltrates (focal, multifocal, and diffuse); and granuloma content (e.g., necrotizing, and non-necrotizing, shown in Supplemental Figure 3 at higher magnification). Additional lesions included fibrin thrombosis with alveolar septal necrosis; cavities with peripheral fibrosis; foamy and multinucleated macrophages with cholesterol clefts; formation of secondary lymphoid follicles; alveolar septal fibrosis; and intra- and extracellular *M. tuberculosis* bacilli described elsewhere [19, 24, 60, 68-71]. Since granuloma necrosis is a key feature of pulmonary TB in humans, we focused on this, and used our automated image analysis method to quantify the ratio of granuloma necrosis in lung tissue sections [57].

**Figure 2.**
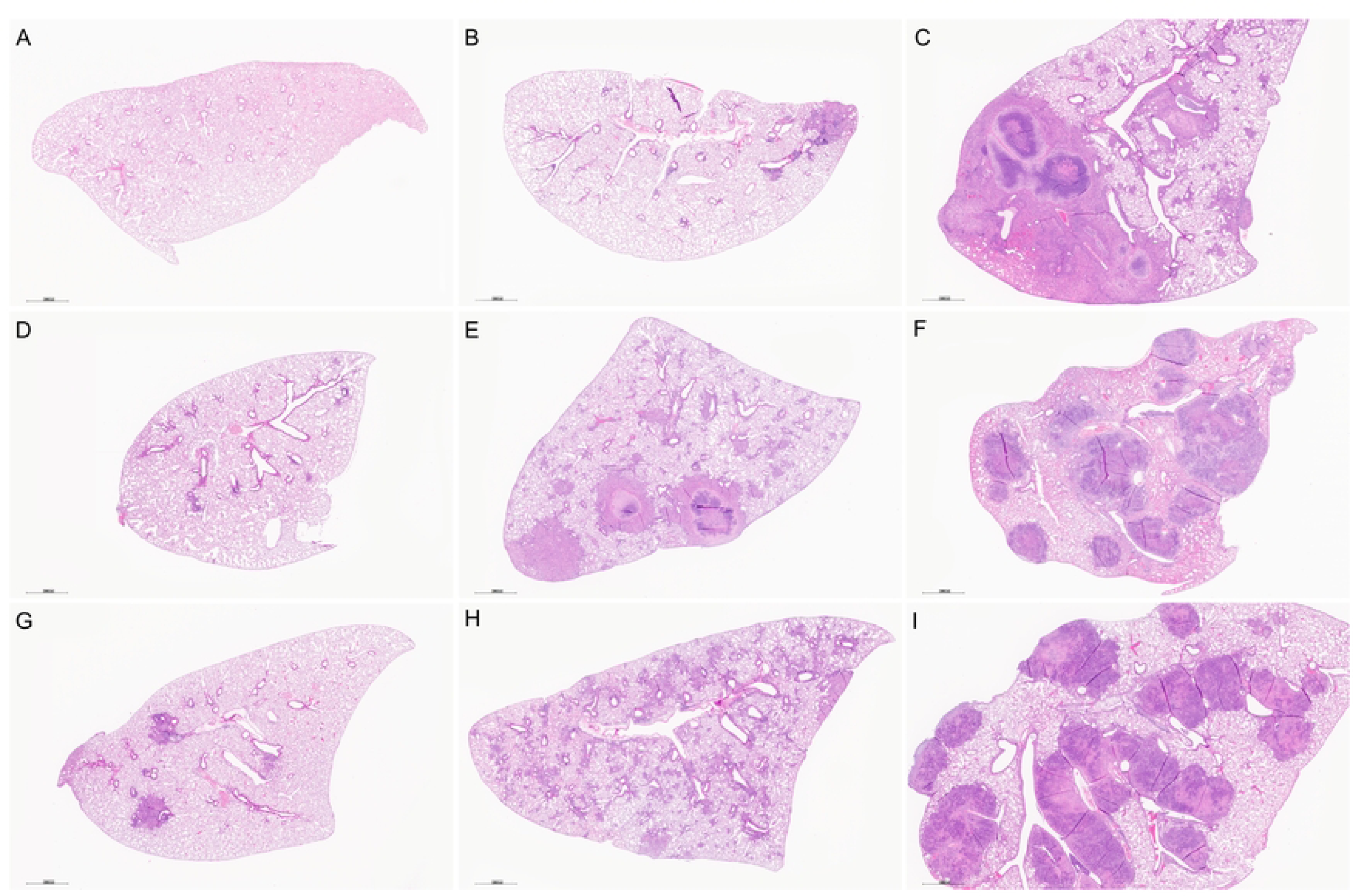
Diversity Outbred (DO) mice develop a spectrum of histopathological lung due to *M. tuberculosis* infection. Lung lobes were formalin-fixed, paraffin-embedded, sectioned, and stained with hematoxylin & eosin. Panel A: Lung section from a non-infected DO mouse. Panels B through I: Lung sections from *M. tuberculosis-*infected DO mice euthanized eight weeks after infection show a spectrum of lesions from mild to severe (upper left to bottom right); focal lesions (e.g., Panel B) to diffuse infiltration (Panel H); and include necrotizing (Panels C, E, F, I) and non-necrotizing granulomas (Panels B, D, G, H). Low magnification (15X).

### Quantification of lung traits: M. tuberculosis burden and immune responses

We quantified *M. tuberculosis* lung burden by counting colonies from lung tissue homogenates and used the remainder lung homogenates for quantification of lung cytokines and chemokines by ELISA [24, 39]. Most lung traits were significantly higher in mice infected with *M. tuberculosis* compared to non-infected mice (Figures 3A), including neutrophil and monocyte/macrophage chemokines (CXCL1,

**Figure 3.**
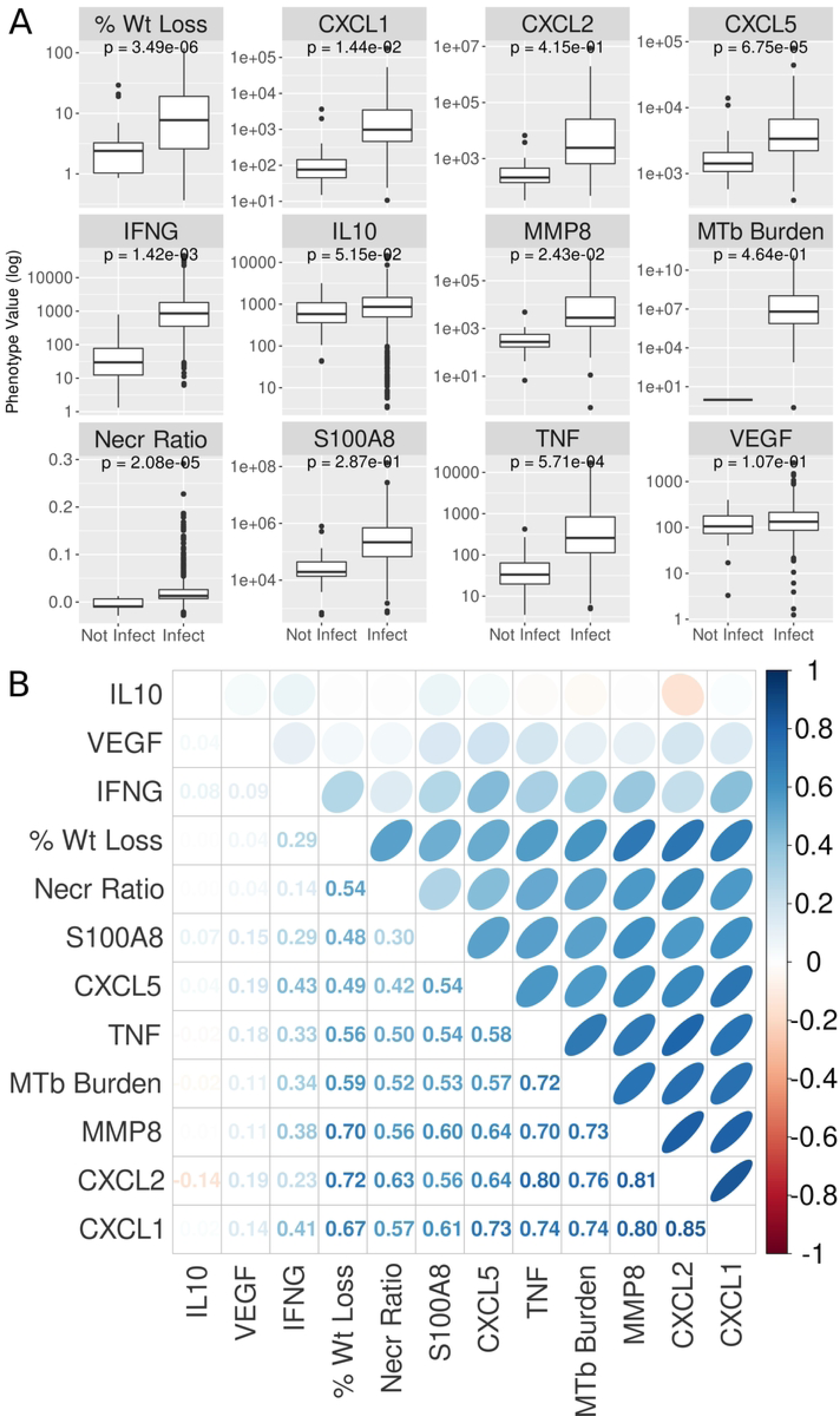
Weight loss, granuloma necrosis, and lung cytokines/chemokines are induced by *M. tuberculosis-*infection of Diversity Outbred (DO) mice and correlate with each other. (A) Traits in *M. tuberculosis-*infected DO mice are higher, with much wider ranges, compared to uninfected DO mice. Each panel shows a boxplot for one phenotype, separated by infection status. Boxes represent the inter-quartile range; center bar is the median and whiskers cover 90% of the data. T-test p-values are shown in each plot. (B) Many traits measured in *M. tuberculosis*-infected DO mice are positively correlated with each other. The lower triangle shows Pearson correlation of pairs of phenotypes. The upper triangle shows these correlations as ellipses, with narrower ellipses indicating higher correlation. All values are colored based on the scale to the left of the plot. Traits are hierarchically clustered on both axes.

CXCL2, CXCL5); mediators of innate immunity (S100A8, Tumor Necrosis Factor (TNF), interleukin (IL)- 10, matrix metalloproteinase-8 (MMP8); mediators of acquired immunity (interferon-gamma (IFN-γ)) and *M. tuberculosis* burden. Pairwise Pearson correlation of all traits in infected mice showed that all the lung traits except IL10 and VEGF positively correlated with each other (Figure 3B). Like previous findings in a small study of Diversity Outbred mice [24], correlations were strongest between *M. tuberculosis* lung burden and mediators of acute neutrophilic inflammation, innate immunity, and extracellular matrix degradation: CXCL1, CXCL2, TNF, and MMP8 with a mean correlation of 0.75.

### Overview of genetic mapping and gene prioritization within QTLs

Figure 5 shows a flow diagram of the types of input data for genetic mapping to identify QTLs, and the subsequent methods of gene prioritization. Briefly, we performed linkage mapping on each trait by regressing each on the additive founder allele dosage at each locus using the R package qtl2 [62]. We selected peaks with a permutation-derived significance threshold of 7.62 (p_GW_ ≤ 0.05) and found seven peaks associated with multiple traits on chromosomes 1, 2, 3, 4, 16, and 17 (Table 1 and Figure 4). We observed that correlated traits colocalized to shared QTLs, and had similar patterns of allele effects, so we used a two-step procedure in which we recorded the confidence interval for peaks with a LOD ≥ 7.62 and then looked for peaks of colocalized traits with a LOD ≥ 6.0 (p_GW_ ≤ 0.6). We reasoned that once we had found the first significant peak for one trait, the threshold for colocalized peaks with the same pattern of founder allele effects should be lower, allowing refinement of the loci.

**Figure 4.**
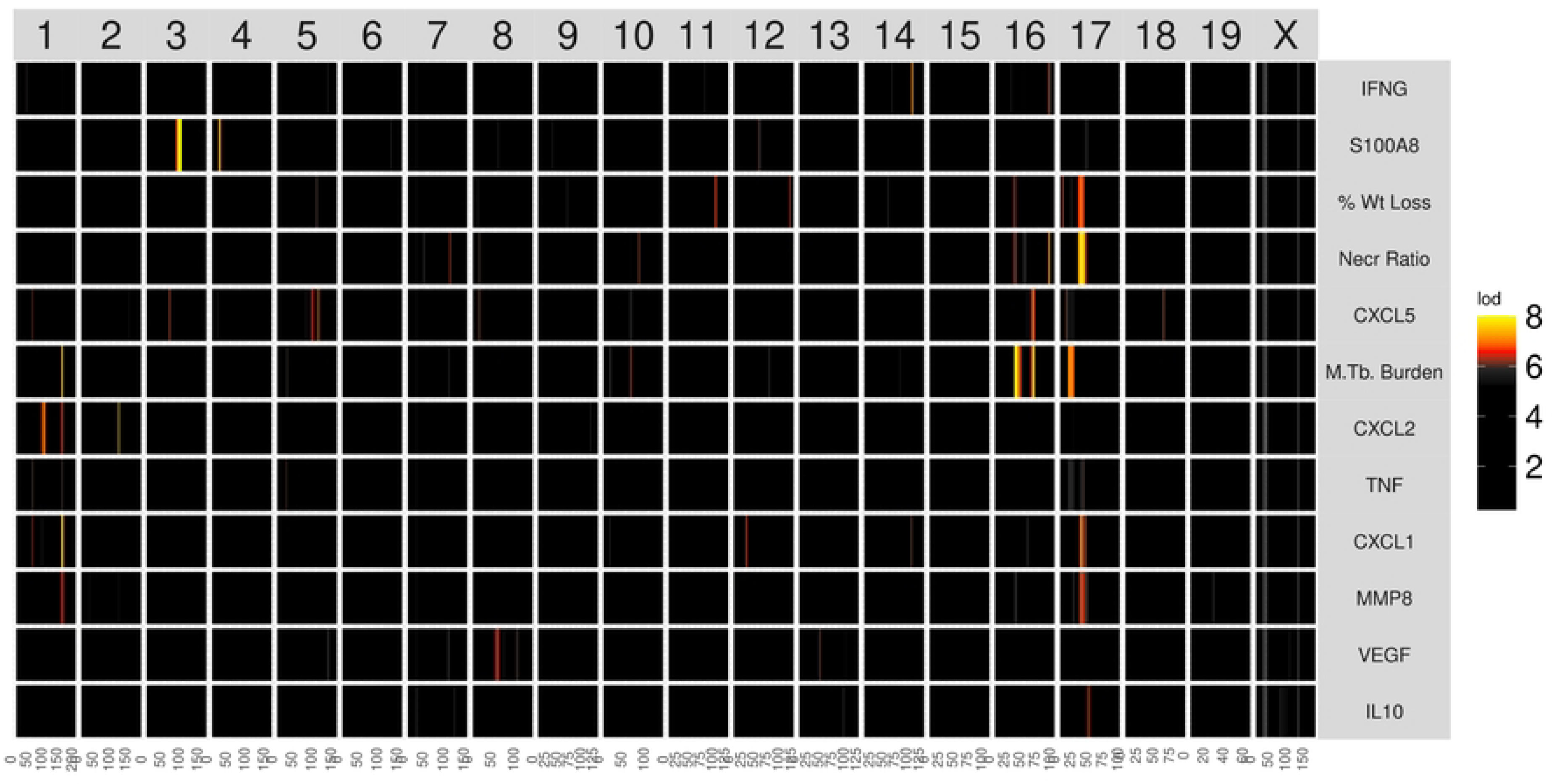
Heatmap of linkage mapping peaks shows patterns of common genetic regulation. The mouse genome, from chromosome 1 through X, is shown on the horizontal axis. Phenotypes are shown on the vertical axis. Each cell shows the LOD score on one chromosome for the phenotype listed on the left, colored by the color scale. The phenotypes are hierarchically clustered based on the correlation between LOD curves, i.e., phenotypes with similar LOD curves are clustered next to each other.

**Figure 5.**
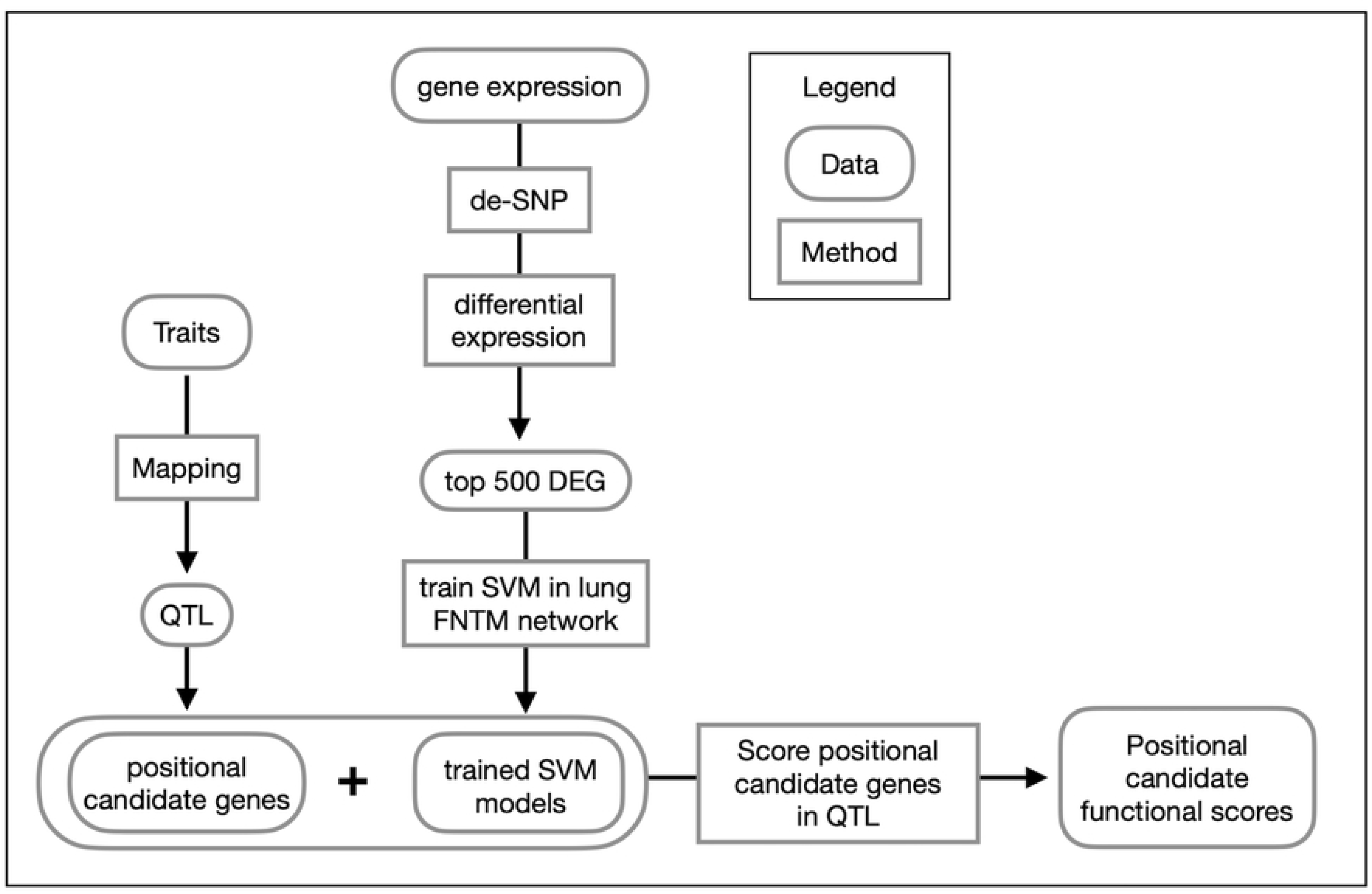
Overview of gene prioritization methods. Traits were mapped to identify positional candidate genes in QTLs. Gene expression data were analyzed for differential gene expression. The top 500 differentially expressed genes (DEG) were used to train SVMs to distinguish these trait-related genes from other genes in the genome using the FNTM mouse lung network. The fitted models were used to score positional candidates in each trait QTL. Positional candidates were then ranked as trait-related based on their functional scores.

### Chromosome 1: Diversity Outbred Tuberculosis Susceptibility locus 1 (Dots1)

*Dots1* is a new QTL on chromosome 1 with a peak LOD >7.6 (p_GW_ < 0.05) at 155.36 M and interval 154.25-156.71 Mb shared by two correlated traits, *M. tuberculosis* burden and CXCL1 (Table 1). Two correlated traits (CXCL2 and MMP8) had lower threshold LODs ≥ 6.0 (p_GW_ ≤ 0.6) mapped to the same position (Figure 4). Notably, these four correlated traits shared patterns of founder allele effects (Supplemental Figure 5), suggesting this QTL contains an important mechanism of genetic regulation for neutrophil-mediated activities, extracellular matrix remodeling, and *M. tuberculosis* growth. To refine the locus, we calculated the first principal component of those four traits and plotted the LOD curve, which also peaked between 154-156 Mb (Figure 6A) and plotted the founder allele effects. A/J, C57BL/6J and WSB/EiJ alleles contributed to higher values of principal component 1 and CAST/EiJ alleles contributed to low allele effects (Figure 6B). We next imputed the founder SNPs onto the Diversity Outbred genomes and performed association mapping in a 10 Mb region around the peak (Figure 6C). Interestingly, the SNPs with highest LOD scores were outside of the peak, and none of the SNPs with the highest LOD scores were missense, stop, or splice site SNPs. This suggested the SNPs in the confidence interval could regulate expression of nearby genes, including some of the 47 protein-coding genes in the interval (Figure 6D and Supplemental File 1). To find and prioritize trait-related gene candidates within *Dots1*, we used the trained SVM model to rank gene candidates based on the strength of their functional relationship in gene expression network modules (Supplemental File 2). *Fam20b* and *Ncf2* ranked highest by functional scoring (Figure 6E). Table 2 summarizes the known annotations, allele effects, founder alleles containing SNPs, and predicted effects of missense SNPs in *Fam20b* and *Ncf2* genes on protein functions from publicly available databases.

**Figure 6.**
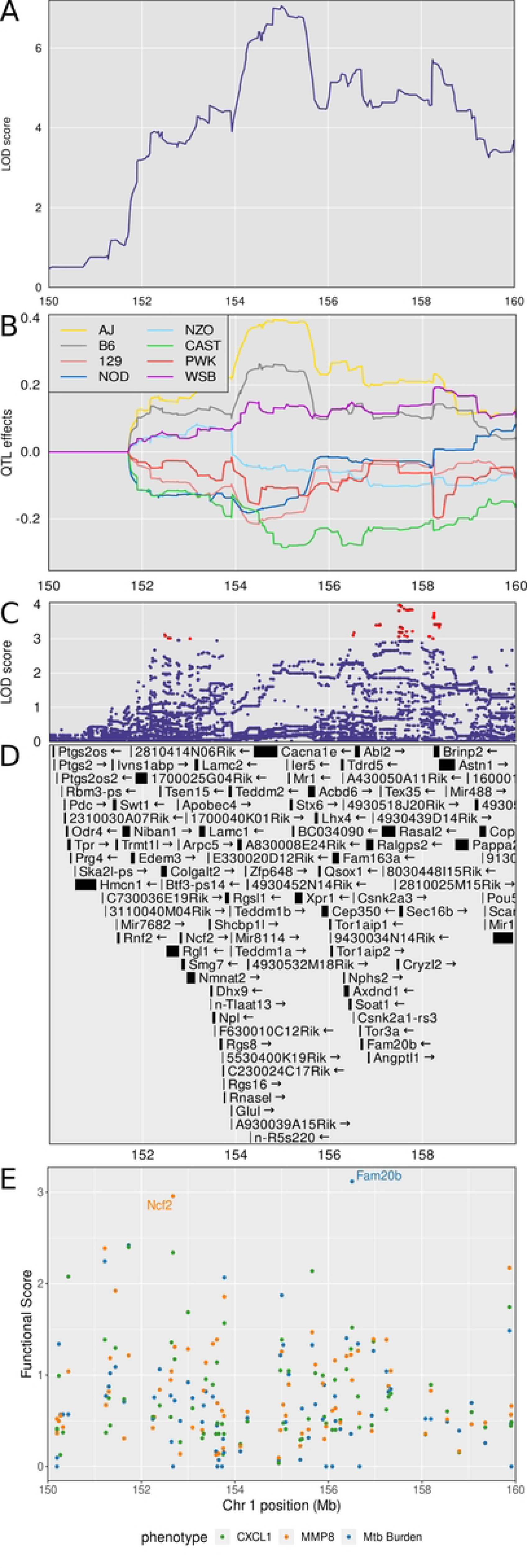
Quantitative Trait Locus (QTL) mapping results of first principal component (PC1) of CXCL1, CXCL2, *M. tuberculosis* burden, and MMP8 identifies *Dots1* on chromosome 1, containing the gene candidates *Fam20b* and *Ncf2*. Panel A shows the LOD curve for PC1 between 150 and 160 Mb on chromosome 1 with peak near 155.36 Mb. Panel B shows the founder allele effects for PC1 in the same genomic interval. Each colored line is the best linear unbiased predictor for one of the founder alleles. Founder colors are shown in the upper left. Panel C shows the LOD score of the imputed SNPs in the same genomic interval. Each point represents the LOD score of one imputed SNP. Panel D shows the genes in the confidence interval. Panel E shows the functional scores for genes in the chromosome 1 QTL. Each dot represents a single gene. Its position on the x axis is its position within the QTL. Its position on the y axis is the functional −log10(UPPR) derived from the SVM. Points are colored based on which trait the functional score corresponds to - green with CXCL1, orange with MMP8, and blue with *M. tuberculosis* burden. *Fam20b* and *Ncf2* genes had the highest functional scores.

### Chromosome 2: Diversity Outbred Tuberculosis Susceptibility locus 2 (Dots2)

*Dots2* is a new QTL, not shared by correlated traits (Table 1), and has a peak LOD of 7.69 (p_GW_ < 0.05) at 22.43 Mb that was associated with lung CXCL2 protein levels (Figure 4). *Dots2* contains 19 protein coding genes (Supplemental File 1). Because this QTL was associated with only 1 trait, gene prioritization by functional scoring was not pursued.

### Chromosome 3: Diversity Outbred Tuberculosis Susceptibility locus 3 (Dots3)

*Dots3* is not a new QTL (Table 1) and overlaps with *tbs1*, a QTL previously identified by crossing A/Sn and I/St inbred mouse strains [72]. These strains are not founder strains of the Diversity Outbred population. *Dots3* was identified by a single peak with a high LOD of 16.57 at 90.69 Mb and interval 90.52-92.02 (p_GW_ < 108) associated with lung S100A8 (calgranulin A) protein levels (Figure 4 and Figure 7A). CAST/EiJ alleles effects were high and PWK/PhJ allele effects were low (Figure 7B). SNPs with the highest LOD scores within the peak are shown (Figure 7C). The interval contains 12 protein coding genes (Supplemental File 1) including the *S100a8* gene (Figure 7D), suggesting that genetic variants which affect *S100a8* transcription regulate S100A8 (calgranulin A) protein levels in *M. tuberculosis* infection. Further, based on based on the strength of the functional relationship in gene expression network modules (Supplemental File 2) the trained SVMs identified *S100a8* as the gene with the highest functional score in *Dots3* (Figure 7E). *Dots3* also contains the gene *S100a9*, which encodes S100A9 (calgranulin B), a protein binding partner of S100A8 (calgranulin A) required to form the heterodimer, calprotectin. Table 2 summarizes the known annotations, allele effects, founder alleles containing SNPs, and predicted effects of missense SNPs in *S100a8* and *S100a9* genes on protein functions from publicly available databases.

**Figure 7.**
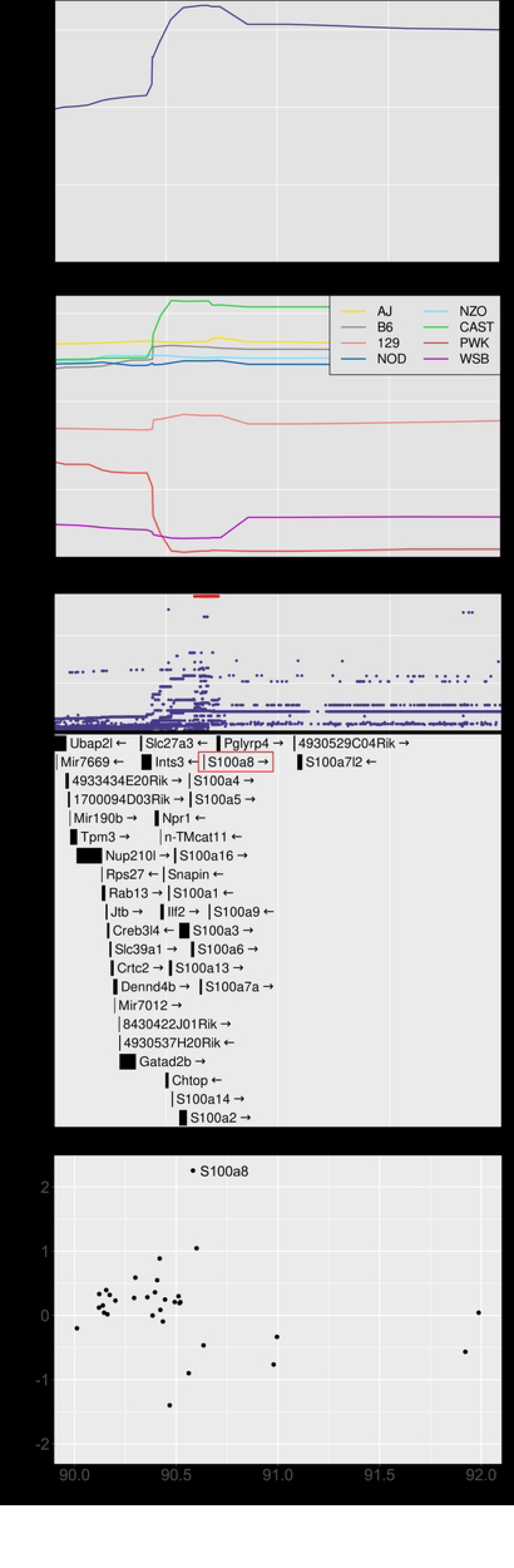
Quantitative Trait Locus (QTL) mapping of lung S100A8 identifies *Dots3* on chromosome 3, containing the gene candidates *S100a8* and *S100a9*. Panel A shows the LOD score in the confidence interval from 90 to 92 Mb on chromosome 3. Panel B shows the founder allele effects within the confidence interval. Panel C shows the LOD score of the imputed SNPs with the highest SNPs colored in red. Panel D shows the gene in the same interval. The gene *S100a8* is directly under the SNPs with the highest LOD scores. Panel E shows the functional scores for genes in the chromosome 1 QTL. Each dot represents a single gene. Its position on the x axis is its position within the QTL. Its position on the y axis is the functional −log10(UPPR) derived from the SVM. The gene *S100a8* had the highest functional score.

### Chromosome 4: Diversity Outbred Tuberculosis Susceptibility locus (Dots4)

*Dots4* is new QTL, not shared by correlated traits (Table 1), and has a peak LOD of 7.64 (p_GW_ < 0.05) at 22.43 Mb and interval 22.18 23.79 Mb associated with lung S100A8 (calgranulin A) protein levels (Figure 4). *Dots4* contains 2 protein coding genes (Supplemental File 1). Because this QTL was associated with only one trait and contained few protein coding genes, prioritization by functional scoring was not pursued.

### Chromosome 16: Diversity Outbred Tuberculosis Susceptibility locus (Dots5)

*Dots5* is a new QTL on chromosome 16, shared by three correlated traits: lung *M. tuberculosis* burden (LOD = 8.45, p_GW_ ≤ 0.01), weight loss, and granuloma necrosis with a peak at 38.3 Mb and interval 33.28-43.28 Mb (Table 1 and Figure 4). We calculated the first principal component of these traits and plotted the LOD curve showing its peak (Figure 8A). The founder allele effects indicate that C57BL/6J alleles contribute to higher values of principal component 1 and that PWK/PhJ and NZO/HILtJ alleles contribute to low effects (Figure 8B). We imputed the founder SNPs onto the Diversity Outbred genomes and performed association mapping around the peak, showing the SNPs with the highest LOD scores (Figure 8C). The SNPs with high LOD scores (Figure 8D) were not missense, stop, or splice site SNPs in the 75 protein coding genes within the interval (Supplemental File 1). By prioritizing genes based on functional relationships in network modules, we identified *Fstl1* and *Itgb5* as functional candidates associated with weight loss, and *Zbtb20* as a functional candidate associated with *M. tuberculosis* burden (Figure 8E, Supplemental File 2). Table 2 summarizes the known annotations, allele effects, founder alleles containing SNPs, and predicted effects of missense SNPs in *Fstl1*, *Itgb5,* and *Zbtb20* genes on protein functions from publicly available databases.

**Figure 8.**
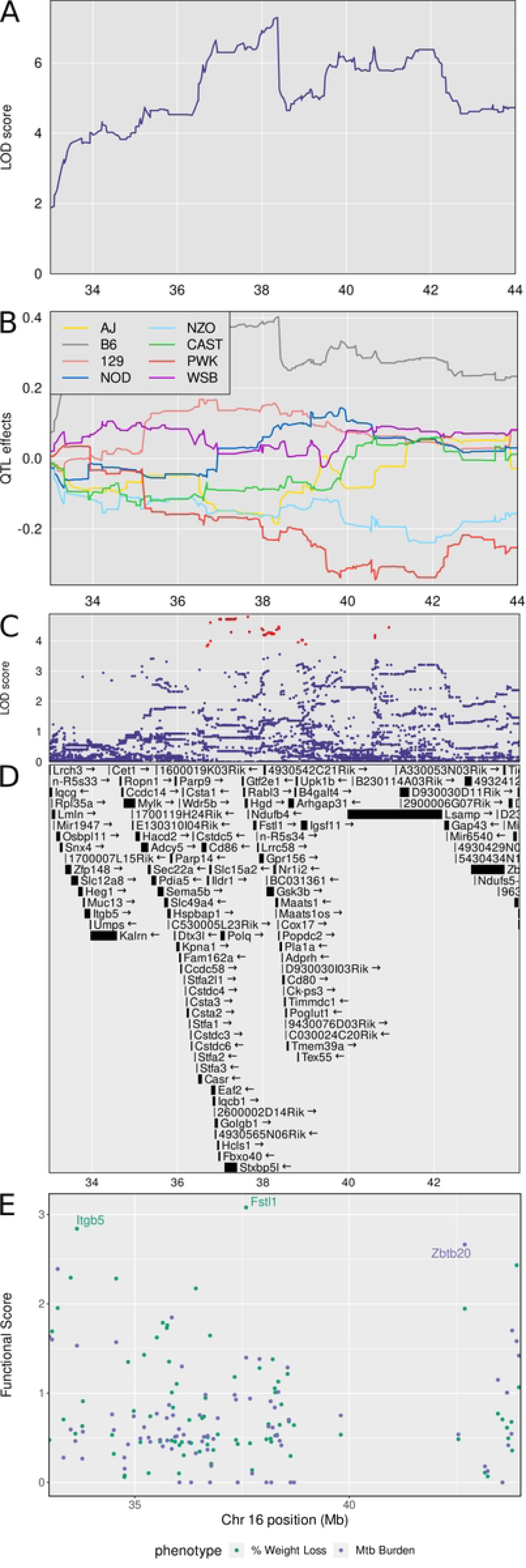
Quantitative Trait Locus (QTL) mapping results of first principal component (PC1) of *M. tuberculosis* burden, weight loss, and granuloma necrosis identifies *Dots5* on chromosome 16, containing gene candidates *Fstl1, Itgb5*, and *Zbtb20*. *M. tuberculosis* lung burden, weight loss, and granuloma necrosis map to a region on chromosome 16 near 38 Mb. Panel A shows the LOD profile for *M.* tuberculosis in the confidence interval. Genomic position on chromosome 16 is on the horizontal axis and the LOD score is on the vertical axis. Panel B shows the founder allele effect in the confidence interval. The vertical axis shows the estimates effect of gaining one founder allele. Panel C shows the SNP LOD score for association mapping using imputed SNPs. Panel D shows the genes in the confidence interval. Panel E shows the functional scores for genes in the chromosome 16 QTL. Each dot represents a single gene. Its position on the x axis is its position within the QTL. Its position on the y axis is the functional −log10(UPPR) derived from the SVM. Points are colored based on which trait the functional score corresponds to - green with weight loss and blue with *M. tuberculosis* burden. *Fstl1* was the top ranked gene overall followed by *Itgb5* and *Zbtb20*.

### Chromosome 16: Diversity Outbred Tuberculosis Susceptibility locus (Dots6)

*Dots6* is new QTL on chromosome 16 and shared by two correlated traits: *M. tuberculosis* burden and CXCL5 with a peak at 52.23 Mb and interval 37.97-57.67 Mb (Table 1 and Figure 4). The interval contains 101 protein coding genes (Supplemental File 1). Because the LOD score was lower than LOD threshold 7.64 for significance (p_GW_ < 0.05), prioritization by functional scoring was not pursued.

### Chr 17: Diversity Outbred Tuberculosis Susceptibility locus (Dots7)

*Dots7* on chromosome 17 is not new, overlaps with *sst5* and *sst6*, QTLs that were previously identified by crossing C3HeB/FeJ and C57BL/6J inbred mouse strains [73]. These strains are not founder strains of the Diversity Outbred population. *Dots7* is associated with *M. tuberculosis* burden and the LOD peaks ∼20 Mb (Table 1 and Figure 4). The interval contains 198 protein coding genes (Supplemental File 1). Because the LOD score was lower than LOD threshold 7.64 for significance (p_GW_ < 0.05), prioritization by functional scoring was not pursued.

### Chr 17: Diversity Outbred Tuberculosis Susceptibility locus (Dots8)

*Dots8* is not a new QTL and also overlaps with *sst5* and *sst6*, two QTLs that were previously identified by crossing C3HeB/FeJ and C57BL/6J inbred mouse strains [73]. Five traits: lung granuloma necrosis (“Necr Ratio”), weight loss, MMP8, CXCL1, and IL-10 mapped to *Dots8.* Lung granuloma necrosis had the highest LOD score (LOD = 8.12, p_GW_ ≤ 0.02) at 35.02 Mb and interval 33.94-41.06 Mb Of those five traits, four positively correlated with each other (Figure 3) and had similar patterns of allele effects, while one associated trait, IL-10 had weak correlations and different patterns of founder allele effects.

We calculated the first principal component of the correlated traits and again performed QTL mapping. Principal component 1 mapped to a wide interval approximately 30-50 Mb with a peak near 38 Mb (Figure 9A). The founder allele effects show PWK/PhJ alleles contribute to high trait values, and NZO/HILtJ and NOD/ShiLtJ alleles contribute to lower values (Figure 9B). We expected to find polymorphisms in the proximal peak of *Dots8* at 34-38 Mb because it contains the mouse histocompatibility-2 (H-2; or Major Histocompatibility Complex-II MHCII). This locus contains many immune response genes known to regulate innate and adaptive immunity and is known to be highly polymorphic. Indeed, the highest SNP association mapping LOD scores were over the mouse H-2 locus, located approximately 36-38 Mb (Figure 9C), and there were 27 SNPs with protein-coding or splice site variation which occurred in 15 genes (Supplemental File 3). Among these were several histocompatibility genes (*H2-M1, H2-M5, H2-M9, H2-M11*) and several tripartite motif (TRIM) family genes (*Trim10, Trim26, Trim31, Trim40*).

**Figure 9.**
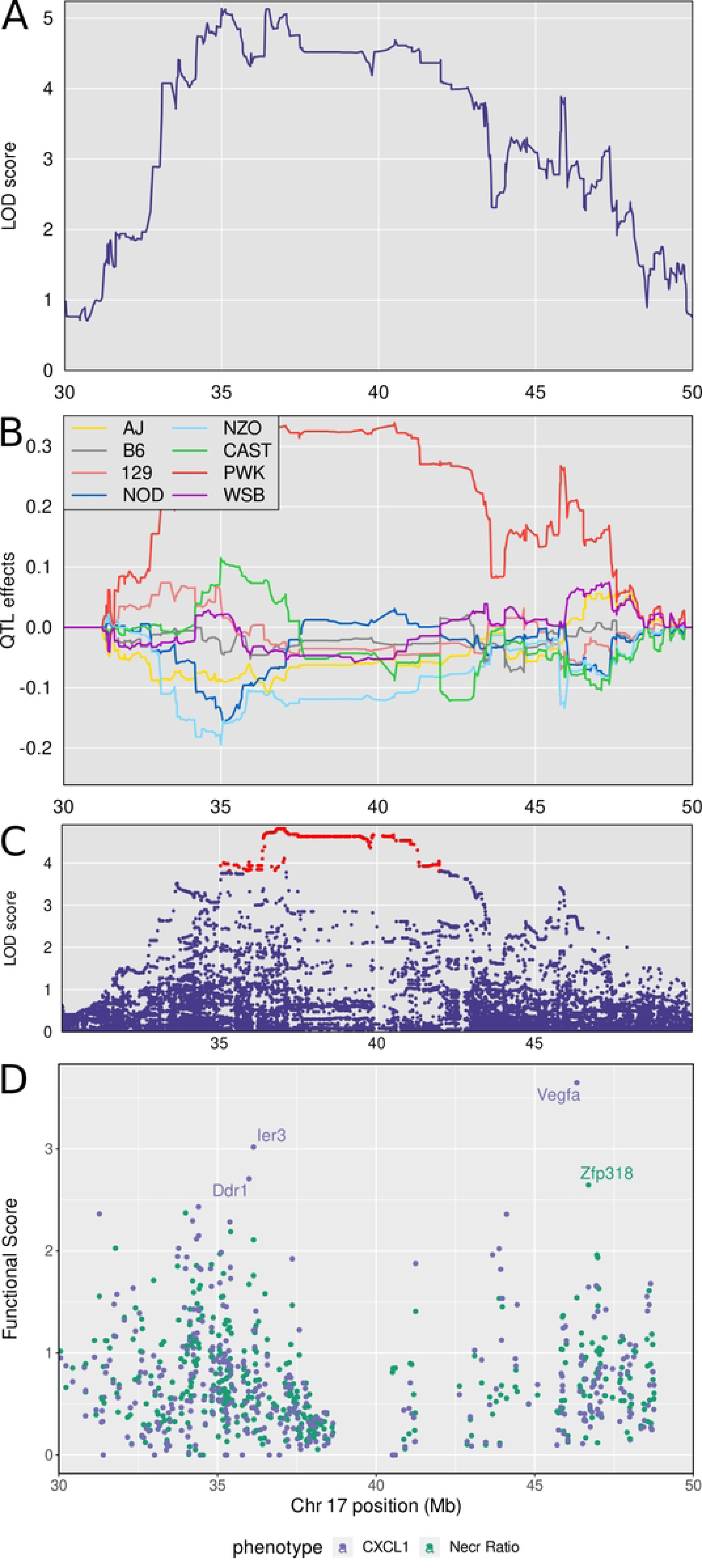
Quantitative Trait Locus (QTL) mapping results of first principal component (PC1) of granuloma necrosis, *M. tuberculosis* burden, weight loss, CXCL1 and MMP8 identifies *Dots7* on chromosome 17, which contains gene candidates *Ier3*, *Ddr1*, and *Zpf318*. Panel A shows the LOD score for PC1 of lung granuloma necrosis ratio, *M. tuberculosis* burden, MMP8, CXCL1, and weight loss in the interval where the phenotypes map. Panel B shows the founder allele effects for the two peaks. Panel C shows the LOD score of the imputed SNPs in the interval, with the highest scoring SNPs colored in red. Panel D shows the locations of the genes in the interval. Panel E shows the functional scores for genes in the chromosome 17 QTL. Each dot represents a single gene. Its position on the x axis is its position within the QTL. Its position on the y axis is the functional −log10(UPPR) derived from the SVM. Points are colored based on which trait the functional score corresponds to - blue with CXCL1 and green with granuloma necrosis. *Vegfa* was the top ranked gene overall followed by *Ier3*, *Ddr1*, and *Zpf318*.

The genes within the broad interval of the *Dots8* are difficult to summarize and interpret as there were 361 protein-coding genes within the 30-50 Mb locus (Supplemental File 1). We prioritized these positional candidate genes again based on their functional relationships in network modules to the correlated traits (Supplemental File 2). This identified candidates *Ddr1, Ier3*, and *Vegfa* associated with CXCL1; and *Zfp318* associated with granuloma necrosis (Figure 9E). Table 2 summarizes the annotations, allele effects, founder alleles containing SNPs, and predicted effects of missense SNPs in *Ddr1, Ier3*, *Vegfa, and Zfp318* genes on protein functions from publicly available databases.

### Selected methodological and gene candidate validation

We performed three different types of validation shown in Supplementary Figure 6. This included (i) survival analysis of Diversity Outbred mice carrying PWK/PhJ alleles at the H-2 locus in *Dots8* on chromosome 17; (ii) quantification of S100A8 protein levels in lungs of *M. tuberculosis* infected PWK/PhJ and CAST/EiJ inbred founder strains; and (iii) infection of gene deficient mice. Notably, infected Diversity Outbred mice carrying at least one copy of the PWK/PhJ allele at the mouse H-2 locus had shorter survival than mice carrying other alleles at the H-2 locus (Supplemental Figure 6A). The lungs of *M. tuberculosis* infected CAST/EiJ inbred mice contained significantly higher levels of S100A8 protein (calgranulin A) than PWK/PhJ inbred mice (Supplemental Figure 6B), confirming the founder allele effects on chromosome 3 *Dots3* QTL, and the levels of S100A8 (calgranulin A) appeared unrelated to *M. tuberculosis* control (Supplemental Figure 6D). Finally, to test *in vivo* effects of one gene candidate, we selected the candidate with the highest LOD score (*S100a8* in *Dots3* QTL on chromosome 3) and obtained C57BL/6 breeding pairs to generate knockout, heterozygous, and wild-type mice. Genotype-tested littermates with null mutation (“knockout”), heterozygous, and wild-type C57BL/6 *S100a8* alleles were infected with *M. tuberculosis*. The absence of one or both copies of the C57BL/6 *S100a8* allele (which is the reference genotype) had minimal impact on *M. tuberculosis* lung burden (Supplemental Figure 6C) suggesting other mechanisms compensate for its absence on the C57BL/6 background.

## Discussion

TB remains a major public health concern in the United States and across the globe, with an estimated 2 billion people infected with *M. tuberculosis*; 8-9 million patients diagnosed each year, and 1-1.5 million deaths annually [2]. Fortunately, most humans (∼90%) are highly resistant to *M. tuberculosis* and clear or control infection [11, 74]. In susceptible adults, active pulmonary TB develops a few years following exposure and tends to occur in young to middle-aged adults in the prime years of their lives [75]. The disease is usually restricted to the lungs and is characterized by granuloma necrosis and cavitation, neutrophilic infiltration, and cachexia [76, 77]. The genetic basis of pulmonary TB is complex and not attributable to single-gene defects that cause severe immune deficiency (i.e., Mendelian susceptibility to mycobacterial disease does not explain pulmonary TB) [15, 78-83]. Although genome-wide association studies have identified loci, gene candidates, and SNPs associated with increased or decreased odds ratios for pulmonary TB, only a few (e.g., Ipr1/SP110b and HLA variants/I-A Major Histocompatibility genes) have been validated [16, 84-90]. This has led investigators to seek alternative experimental mouse models such as Diversity Outbred mice and Collaborative Cross recombinant inbred strains to examine effects of genetics on host responses to *M. tuberculosis* [24-26, 38].

An advantage of the Diversity Outbred mouse population is that infection with *M. tuberculosis* induces phenotypes that are rare in common laboratory inbred strains of mice [15-17]. Further, a growing body of evidence shows similarities in *M. tuberculosis-*infected Diversity Outbred mice and humans in biomarkers, gene expression signatures, and BCG vaccination [38, 39, 67, 68, 91]. The phenotypic similarities suggest that humans and Diversity Outbred mice may share underlying genetic pathways of immunity and disease. And, because SNP variants in the Diversity Outbred mouse genomes are dense, with balanced allele frequencies, any gene that plays a role in disease is theoretically detectable [92]. This eliminates a problem common to human genetic studies where under-represented alleles cannot be confidently associated with disease phenotypes because they are low-frequency genetic events.

We performed genetic mapping in *M. tuberculosis-*infected Diversity Outbred mice and used orthogonal methods to rationally select candidate genes. We first used DOQTL mapping, which relies entirely on phenotypic and genetic variation, to find eight QTLs on six different chromosomes named *Dots1* through *Dots8*. To refine loci, we then subjected the QTLs on chromosomes 1, 16, and 17 (*Dots1, Dots5, and Dots* 8) by mapping the first principal component of the correlated traits with similar patterns of allele effects that colocalized to the same interval. Finally, we applied a gene-based machine-learning SVM to identify and rank gene candidates based on functional scores. The sequential methods narrowed the candidate gene list to eleven polymorphic, protein coding genes. Finally, the SNPs were critically examined using publicly available databases to find four candidates (*S100a8, Itgb5, Fstl1, Zfp318*) where missense SNPs are predicted to have deleterious effects on protein function.

All eleven candidates have roles in infection, inflammation, cell migration, extracellular matrix remodeling, or intracellular signaling. Of those, only one (mouse *Ncf2* in *Dots1* on chromosome 1) has a human homologue where a single SNP (G nucleotide in human NCF2 rs10911362) may provide a protective effect it lowered the odds ratio of pulmonary TB [93]. Absence of *Ncf2* on the C57BL/6 inbred mice this temporarily impairs resistance to *M. tuberculosis* infection by abrogating superoxide production, but the defect does not affect overall survival due to compensation by T cell mediated immunity [94]. Experimental validation of mouse *Ncf2* and human NCF2 polymorphisms remains to be confirmed.

We identified *Fam20b* in *Dots1* QTL on chromosome 1. The gene encodes a xylosylkinase that functions in glycosaminoglycan synthesis to produce extracellular matrix components in tissues. Deficiencies are embryologically lethal or cause cranioskeletal malformations [44, 45, 95, 96]. A functionally homologous enzyme phosphorylates cadherins [97] which regulate immune cell migration [98] by interacting with extracellular matrix components, and since cell migration is required to form mycobacterial granulomas, *Fam20b* polymorphisms may alter host susceptibility to *M. tuberculosis* by changing extracellular matrix.

We identified *S100a8* and *S100a9* in *Dots3* QTL on chromosome 3, which encode S100A8 (calgranulin A) and S100A9 (calgranulin B). The proteins form monomers, homodimers, heterodimers, and multimers in inflammation, host defense, and nociception [99-102]. Some forms activate Toll-Like receptor 4 signaling; some activate the receptor for advanced glycation end-products; and some sequester calcium, zinc, and manganese metal ions [99, 103, 104]. In pulmonary TB, S100A9 contributes to neutrophil localization to granulomas, and both S100A8 and S100A9 are protein biomarkers of TB- related lung damage [38, 39, 104-107]. Interestingly, 4 polymorphisms in *S100a8* were predicted to have deleterious effects on function but the lack of S100a8 did not change the ability of C57BL/6 inbred mice to restrict *M. tuberculosis* growth. Further investigation would be required to determine the effects on other host outcomes.

We identified *Itgb5* in *Dots5* QTL on chromosome 16 as a gene candidate. *Itgb5* encodes the beta 5 (β5) integrin subunit which dimerizes with the alpha v subunit to mediate cell adhesion and signaling by binding to fibronectin and vitronectin [41]. Notably, the β5 subunit is on the surface of cancer cells, and normal epithelial cells and activated endothelial cells but not on lymphoid or myeloid cells [41, 108- 114]. To our knowledge, neither the mouse nor human gene, nor subunit β5, nor the αvβ5 integrin heterodimer have been deeply investigated in pulmonary TB.

We identified *Fstl1* in *Dots5* QTL on chromosome 16. The primary transcript encodes microRNA (miR)-198. The product is a secreted glycoprotein, FSTL1 with activities in angiogenesis, cell proliferation, differentiation, embryogenesis, metastasis, and wound healing; specifically reducing inflammation and fibrosis in cardiovascular disease [43, 115-118]. Notably, *Fstl1* affects survival of *M. tuberculosis-*infected macrophages [34, 35, 119]. Given the central role of macrophages, inflammation, and fibrosis in *M. tuberculosis* infection, understanding how *Fstl1* polymorphisms and FSTL1 function *in vivo* may inform TB pathogenesis, and possibly targets for host-directed therapy.

We identified *Zbtb20* in *Dots5* QTL on chromosome 16. The gene encodes a transcriptional repressor involved in glucose homeostasis; growth; hematopoiesis; innate immunity; neurogenesis; and B cell development and long-term survival of plasma cells [46-49, 120-124]. Natural mutations occur in humans with Primrose Syndrome, although immune deficiencies are not reported [125]. To our knowledge, there are no studies on *Zbtb20* and *M. tuberculosis* infection or pulmonary TB. However, in *Listeria monocytogenes* infection*, Zbtb20*-deficiency improved CD8 T cell memory functions due to efficient use of diverse fuel sources [49]. Whether the same is true in pulmonary TB is unknown.

We identified *Ddr1* in *Dots8* QTL on chromosome 17. *Ddr1* encodes for the discoidin domain receptor 1 (DDR1), which interacts with collagen [42]. Initial studies suggested DDR1 function was restricted to epithelial cells; however, recent work shows expression on solid tumors, metastatic cells, and mouse histiocytic cancer cell lines, J774 and Raw264.7 [126-132]. DDR1 has additional roles in demyelination, fibrosis, vitiligo, and wound healing, and it is also a promising target for anti-fibrotic therapy [133-137]. Whether *Ddr1* (mouse) or DDR1 (human) gene polymorphisms contribute to pulmonary TB, or whether it could be a target for anti-fibrotic therapy in TB are areas open for investigation.

We identified the immediate early response gene, *Ier3* in *Dots8* QTL on chromosome 17. *Ier3* transcription is triggered by cytokines, hormones, DNA damage, and infections. The protein, IER3, regulates apoptosis, DNA repair, differentiation, and proliferation by interfering with NF-κB, MAPK/ERK and PI3K/Akt signaling pathways [50-52, 138-141]. Mice lacking *Ier3* are more susceptible to *Leishmania* [142], an intracellular pathogen that shares some similar immune responses profiles with those induced by *M. tuberculosis* but we did not find studies showing that mutated *Ier3* also increases susceptibility to *M. tuberculosis*. One *in vitro* study of human macrophages, however, had high levels of IER3 mRNA following infection with a hypervirulent strain of *M. tuberculosis* [143] indicating the transcriptional pathway is triggered.

We identified *Vegfa* in *Dots8* QTL on chromosome 17. Mouse *Vegfa* and human VEGFA, encode for a heparin-binding protein and essential growth factor that induces proliferation, migration, and permeability changes in vascular endothelial cells by binding VEGFR1 and VEGFR2 [144-147]. Roles for VEGF in pathogenesis and diagnostics for extrapulmonary TB, cavitary TB, and active TB have been published [148-150]. Myeloid-specific gene deletion of *Vegfa* extended survival of C57BL/6J inbred mice infected with *M. tuberculosis* [151], highly noteworthy because very few gene deletions improve survival. Whether *Vegfa* or VEGFA polymorphisms have the same effect is unknown.

Lastly, we identified *Zfp318* in *Dots8* QTL on chromosome 17. The gene encodes the transcription factor, zinc finger protein 318 and it is expressed in testes, hematopoietic, and lymph nodes [152]. In B cells, the protein represses transcription required for class switching, helping to maintain B cell anergy and prevent autoimmunity [153-155]. Database and literature searches identified no publications on mouse *Zpf318* or human ZPF318 in infectious diseases.

When we compared genetic mapping results from Diversity Outbred mice to results from colleagues using Collaborative Cross inbred strains [26], QTLs and gene candidates did not overlap although we measured a few of the same traits by standard laboratory methods (e.g., body weight changes, lung *M. tuberculosis* burden, and lung CXCL1 by immunoassays). This suggests that phenotype-genotype relationships in the Collaborative Cross strains may be fundamentally different than Diversity Outbred mouse population (despite sharing the same eight inbred founder strains), possibly because of high levels of heterozygosity in the Diversity Outbred population. Other reasons could be differences in routes of infection that change the host cell types first encountering *M. tuberculosis* bacilli which alters antigen presentation, T-cell, and B-cell priming. Here, we modeled natural aerosol exposure by delivering a low dose of approximately 20-100 bacilli to the lungs of Diversity Outbred mice in nebulizer-delivered aerosol mist, and then focused on quantification of lung disease. In contrast, Smith *et al* [26] took a different approach by using intravenous infection with 1×106 bacilli to take advantage of their rich library of transposon mutants, allowing detailed assessment of pathogen-associated QTLs. As the intravenous route of infection favors rapid induction of acquired immunity by delivering bacilli directly to lymphoid organs (i.e., spleen, thoracic, and abdominal lymph nodes by portal and systemic circulation), this approach maximized identification of unique Host-Interacting-with Pathogen QTLs and resulted in a prioritized list of candidate genes involved in immunity.

Overall, by using a systems genetics approach focused on the lungs, we multiple new and existing QTLs, and 11 candidate genes. Of those, gene products for five (*Ncf2, Fstl1, Zbtb20 Vegfa, Zfp318*) have known roles in recruitment, activation, or regulation of effector functions of immune cells (e.g., neutrophils, monocytes, macrophages and CD8 T cells). The gene products for three candidates (*Fam20b, Itgb5, Ddr1*) have known roles in epithelial cell, endothelial cell, and (possibly) macrophage adhesion to extracellular matrix glycoproteins or are involved in remodeling of extracellular matrix. The gene products for two candidates (*S100a8* and *S100a9*) have complex and context-dependent roles in innate immune response signaling and in host defenses. Finally, the gene product of one candidate (*Ier3*) controls early stress responses of cells, including cell survival and death pathways. Ten of the eleven candidates have annotated polymorphisms; six have missense SNPs in protein coding regions; and the SNPs in four candidates (*S100a8, Itgb5, Fstl1, and Zfp318)* are predicted to have deleterious consequences on protein functions. Together, these results yield a short list of candidates that may be major regulators of host necrotizing and inflammatory responses during *M. tuberculosis* infection and pulmonary TB disease progression. Future studies will focus on testing effects of these gene candidates and polymorphisms *in vivo* and identification of pathogenic molecular and cellular mechanisms.

## Acknowledgements

We thank Ms. Julie Tzipori, Mr. Curtis Rich, Mr. Donald Girouard, and Dr. Sam Telford III for study support at the New England Regional Biosafety Laboratory at Tufts University Cummings School of Veterinary Medicine, North Grafton, MA. NIH NIAID UC6A1066843 supported construction of the New England Regional Biosafety Laboratory. Ms. Frances Brown, Ms. Linda Wrijil, Ms. Sarah Ducat, Ms. Gina Scarglia, and Dr Amanda Martinot provided histology services at Tufts University’s Cummings School of Veterinary Medicine. Whole slide imaging was performed by the Digital Histology Shared Resource at Vanderbilt University Medical Center. The following reagents were obtained through BEI Resources, supported by NIH NIAID: ESAT-6, Recombinant Protein Reference Standard, NR-49424; CFP-10, Recombinant Protein Reference Standard, NR49425; Plasmid pMRLB.7 Containing Gene Rv3875 (Protein ESAT-6) from Mycobacterium tuberculosis, NR-50170; Plasmid pMRLB.46 Containing Gene Rv3874 (Protein Cfp10) from *Mycobacterium tuberculosis*, NR-13297; *Mycobacterium tuberculosis*, Strain H37Rv, Culture Filtrate Proteins, NR-14825; and *Mycobacterium tuberculosis*, Strain H37Rv, Cell Wall Fraction, NR- 14828. All microarray protocols were carried out by the Boston University Microarray and Sequencing Resource (BUMSR) core facility, and we thank Mr. Eduard Drizik of the BUMSR for initial microarray analyses. The following individuals are thanked for their technical expertise with *in vitro* and *in vivo* experiments: Ms. Victoria Mello at Tufts Cummings School of Veterinary Medicine, North Grafton, MA; Mr. Austin Hossfeld at The Ohio State University, Columbus, OH; and Dr. Joanne Turner at Texas Biomedical Research Institute, San Antonio, TX.

## AUTHOR SUMMARY

We investigated the genetic basis of susceptibility to *Mycobacterium tuberculosis* using Diversity Outbred mice, a mouse population suited for studies on complex genotype-phenotype relationships. We identified multiple new genetic loci as well as two previously identified loci. Interestingly, we found three loci associated with multiple disease traits, which indicates genes within the loci are likely major regulators of host inflammatory responses which permit *M. tuberculosis* growth. These three loci contain at least four gene candidates with single nucleotide polymorphisms that are predicted to have deleterious effects upon protein functions.

---

**Supplemental Figure 1. Mouse body weight following a low dose of aerosolized *M. tuberculosis* strain Erdman.** Mice were infected with a low dose of *M. tuberculosis* strain Erdman by aerosol. Body weight of identically housed, age-, gender-, and generation-matched non-infected Diversity Outbred (DO) controls (n = 49) compared to baseline are shown over time (A). Body weight changes of Progressor DO mice (n = 195); Controller DO mice (n = 145); and C57BL/6J inbred founder strain mice that succumbed to pulmonary TB (n = 39), are shown over time compared to pre-infection baseline (B, C, D). All mice were weighed 1 to 3 days prior to *M. tuberculosis* infection, at least twice per week throughout infection, and immediately before euthanasia. Each line is the body weight expressed as a percentage of initial pre-infection body weight.

**Supplemental Figure 2. Clinical correlates of survival due to pulmonary TB in Diversity Outbred (DO) mice following exposure to a low dose of aerosolized *M. tuberculosis* strain Erdman.** Age-, gender-, and generation-matched DO mice were assigned to cages at random, and infected (or not infected) with a low dose of *M. tuberculosis* strain Erdman by aerosol exposure. All mice were initially weighed 1-3 days prior to infection, at least twice per week during infection, and immediately before euthanasia. Panel (A) shows retrospective analysis of initial body weights of Non-infected mice (n = 76) compared to pre-infection body weights of Progressors (n = 298) and pre-infection body weights of Controllers (n = 195), shown as box-and-whisker plots with the line at the mean for each group, and whiskers at the minimum and maximum. Data were analyzed by 1-way ANOVA with Tukey’s multiple comparisons test ***p<0.001; ****p<0.0001. Panel B shows the rate of weight loss (gm/day) and duration of body weight (BW) loss in days were negatively correlated. Panel C shows the duration of BW loss was strongly, positively, and linearly correlated with survival by Spearman correlation analysis (r = 0.848 with dashed lines indicating the 95% confidence interval, 0.8204 to 0.8717, p<0.0001). Panel D is a correlation matrix to show how survival and 8 clinical indicators of pulmonary TB in Diversity Outbred mice correlate with each other. Only correlations with p-values <0.00001 are shown on the matrix. Cells marked by an “X” were not significantly correlated.

**Supplemental Figure 3. Examples of necrotizing and non-necrotizing lesions in *M. tuberculosis* infected Diversity Outbred (DO) mice.** Lung lobes were formalin-fixed, paraffin-embedded, sectioned, and stained with carbol fuschin and counterstained with hematoxylin & eosin. Panels A and B: High magnification images of necrotizing lung lesions. One example contains abundant pyknotic nuclear debris (A) and one example contains abundant fibrin, eosinophilic cellular debris, and less nuclear debris (B). Panels C and D: High magnification images of non-necrotizing lung lesions. Both examples contain predominantly viable cells, including macrophages, foamy macrophages, and foci of lymphocytes (400X).

**Supplemental Figure 4. Common founder allele effects for four traits on distal chromosome 1 QTL.** Founder allele effects of the four phenotypes with genetic mapping peaks having LOD > 6.0 on chromosome 1 at 155.36 Mb. All four phenotypes have similar allele effects. Each panel shows the founder allele effects for the phenotype listed in the title. Founders are on the horizontal axis and the standardized allele effect are on the horizontal axis.

**Supplemental Figure 5. Receiver operator characteristic (ROC) curves for SVM training on traits used in gene prioritization.** Each panel shows the true positive rate of the trained SVM as a function of the false positive rate for each trait. The area under the curve (AUC) is noted for each panel.

**Supplemental Figure 6. Validation of QTL mapping results.** Panel A: *M. tuberculosis* infected Diversity Outbred (DO) mice with one or more PWK/PhJ alleles at the mouse H-2 locus on chromosome 17 in *Dots8* (near 36 Mb) have significantly reduced survival compared to DO mice carrying other alleles (p = 0.00075, Cox-PH test). Kaplan-Meier curves of survival of *M. tuberculosis* infected mice carrying PWK/PhJ allele (red) or any other founder allele (black). Days of survival is shown on the horizontal axis and the proportion of mice surviving is shown on the vertical axis. Panel B: Lungs from *M. tuberculosis* infected CAST/EiJ inbred mice contain significantly more S100A8 protein (calgranulin A) than lungs of PWK/PhJ inbred mice. PWK/PhJ (red) and CAST/EiJ (green) inbred founder strains with 4-6 mice per strain time point, analyzed by Mann-Whitney *t-*tests within each time point, *p<0.05. Panel C: *M. tuberculosis* infected *S100a8* knockout (KO), heterozygotes (HET), wild-type (WT) C57BL/6 inbred mice were euthanized at the time points indicated on the X-axes, and *M. tuberculosis* lung burden assessed by CFUs with total combined 15-22 mice per genotype per time point from 2 independent experiments, shown as average and standard error of the mean. No significant (ns) differences were identified within each time point by mixed effects ANOVA with Tukey’s post-test (p<0.05). Panel D shows *M. tuberculosis* burden in the lungs of PWK/PhJ (red) and CAST/EiJ (green) inbred founder strains with 4-6 mice per strain time point. No significant differences were identified by Mann-Whitney *t-*tests within each time point, although there was a trend for higher bacterial burden at day 20 post infection in lungs from CAST/EiJ inbred mice as compared to PWK/PhJ.

**Supplemental File 1** This file is an Excel workbook containing worksheets that list all protein-coding genes in each QTL with the functional candidates highlighted.

**Supplemental File 2** This is an Excel fie that lists the top 10 functional candidates for each trait in each QTL.

**Supplemental File 3** This is an Excel file that lists genes within chromosome 17 QTL.

## References

1. Organization, W.H. *14.9 million excess deaths associated with the COVID-19 pandemic in 2020 and* 2021. 2022 5/5/2022 5/9/2022]; Available from: https://www.who.int/news/item/05-05-2022-14.9-million-excess-deaths-were-associated-with-the-covid-19-pandemic-in-2020-and-2021.

2. Organization, W.H., *Global tuberculosis report* 2021. 2021, World Health Organization: Geneva, Switzerland. p. 57.

3. WHO, Global Tuberculosis Report: Multidrug-/ rifampicin-resistant TB (MDR/RR-TB):Update 2017. 2017.

4. WHO, Global Tuberculosis Report. 2013.

5. Soleimanpour, S., et al., A century of attempts to develop an effective tuberculosis vaccine: Why they failed? Int Immunopharmacol, 2022. 109: p. 108791.

6. Heidary, M., et al., Tuberculosis challenges: Resistance, co-infection, diagnosis, and treatment. Eur J Microbiol Immunol (Bp), 2022.

7. Kassa, G.M., et al., Predictors of mortality among multidrug-resistant tuberculosis patients in central Ethiopia: a retrospective follow-up study. Epidemiol Infect, 2020. 148: p. e258.

8. Tiemersma, E.W., et al., Natural history of tuberculosis: duration and fatality of untreated pulmonary tuberculosis in HIV negative patients: a systematic review. PLoS One, 2011. 6(4): p. e17601.

9. Vilaplana, C. and P.J. Cardona, The lack of a big picture in tuberculosis: the clinical point of view, the problems of experimental modeling and immunomodulation. The factors we should consider when designing novel treatment strategies. Front Microbiol, 2014. 5: p. 55.

10. Achkar, J.M. and E.R. Jenny-Avital, Incipient and subclinical tuberculosis: defining early disease states in the context of host immune response. J Infect Dis, 2011. 204 Suppl 4: p. S1179–86.

11. Verrall, A.J., et al., Early clearance of Mycobacterium tuberculosis: a new frontier in prevention. Immunology, 2014. 141(4): p. 506–13.

12. Flynn, J.L. and J. Chan, Immunology of tuberculosis. Annual Review of Immunology, 2001. 19: p. 93–129.

13. Hunter, R.L., C. Jagannath, and J.K. Actor, Pathology of postprimary tuberculosis in humans and mice: contradiction of long-held beliefs. Tuberculosis, 2007. 87(4): p. 267–78.

14. Azad, A.K., W. Sadee, and L.S. Schlesinger, Innate immune gene polymorphisms in tuberculosis. Infect Immun, 2012. 80(10): p. 3343–59.

15. Gibbs, K.D., B.H. Schott, and D.C. Ko, The Awesome Power of Human Genetics of Infectious Disease. Annu Rev Genet, 2022.

16. Medhasi, S. and N. Chantratita, Human Leukocyte Antigen (HLA) System: Genetics and Association with Bacterial and Viral Infections. J Immunol Res, 2022. 2022: p. 9710376.

17. Varshney, D., et al., Systematic review and meta-analysis of human Toll-like receptors genetic polymorphisms for susceptibility to tuberculosis infection. Cytokine, 2022. 152: p. 155791.

18. Williams, A. and I.M. Orme, Animal Models of Tuberculosis: An Overview. Microbiol Spectr, 2016. 4(4).

19. Kramnik, I. and G. Beamer, Mouse models of human TB pathology: roles in the analysis of necrosis and the development of host-directed therapies. Semin Immunopathol, 2016. 38(2): p. 221–37.

20. Clark, S., Y. Hall, and A. Williams, Animal models of tuberculosis: Guinea pigs. Cold Spring Harb Perspect Med, 2015. 5(5): p. a018572.

21. Scanga, C.A. and J.L. Flynn, Modeling tuberculosis in nonhuman primates. Cold Spring Harb Perspect Med, 2014. 4(12): p. a018564.

22. Dharmadhikari, A.S. and E.A. Nardell, What animal models teach humans about tuberculosis. American journal of respiratory cell and molecular biology, 2008. 39(5): p. 503–8.

23. Helke, K.L., J.L. Mankowski, and Y.C. Manabe, Animal models of cavitation in pulmonary tuberculosis. Tuberculosis, 2006. 86(5): p. 337–48.

24. Niazi, M.K., et al., Lung necrosis and neutrophils reflect common pathways of susceptibility to Mycobacterium tuberculosis in genetically diverse, immune-competent mice. Dis Model Mech, 2015. 8(9): p. 1141–53.

25. Smith, C., et al., Tuberculosis Susceptibility and Vaccine Protection Are Independently Controlled by Host Genotype. mBio, 2016. 7(5): p. e01516.

26. Smith, C.M., et al., Host-pathogen genetic interactions underlie tuberculosis susceptibility in genetically diverse mice. Elife, 2022. 11.

27. Churchill, G.A., et al., The Diversity Outbred mouse population. Mamm Genome, 2012. 23(9-10): p. 713–8.

28. Tabangin, M.E., J.G. Woo, and L.J. Martin, The effect of minor allele frequency on the likelihood of obtaining false positives. BMC Proc, 2009. 3 **Suppl** 7(Suppl 7): p. S41.

29. Gatti, D.M., et al., Quantitative trait locus mapping methods for diversity outbred mice. G3 (Bethesda), 2014. 4(9): p. 1623–33.

30. Guan, Y., et al., Functional genomics complements quantitative genetics in identifying disease-gene associations. PLoS Comput Biol, 2010. 6(11): p. e1000991.

31. Tyler, A.L., et al., Network-Based Functional Prediction Augments Genetic Association To Predict Candidate Genes for Histamine Hypersensitivity in Mice. G3 (Bethesda), 2019. 9(12): p. 4223–4233.

32. Goya, J., et al., FNTM: a server for predicting functional networks of tissues in mouse. Nucleic Acids Res, 2015. 43(W1): p. W182–7.

33. Crouser, E.D., et al., A Novel In Vitro Human Granuloma Model of Sarcoidosis and Latent Tuberculosis Infection. Am J Respir Cell Mol Biol, 2017. 57(4): p. 487–498.

34. Zhang, Z.M., et al., TLR-4/miRNA-32-5p/FSTL1 signaling regulates mycobacterial survival and inflammatory responses in Mycobacterium tuberculosis-infected macrophages. Exp Cell Res, 2017. 352(2): p. 313–321.

35. Deng, Q., et al., Circ_0001490/miR-579-3p/FSTL1 axis modulates the survival of mycobacteria and the viability, apoptosis and inflammatory response in Mycobacterium tuberculosis-infected macrophages. Tuberculosis (Edinb), 2021. 131: p. 102123.

36. Duan, X.Y., et al., Baicalin attenuates LPS-induced alveolar type II epithelial cell A549 injury by attenuation of the FSTL1 signaling pathway via increasing miR-200b-3p expression. Innate Immun, 2021. 27(4): p. 294–312.

37. Henkel, M., et al., Regulation of Pulmonary Bacterial Immunity by Follistatin-Like Protein 1. Infect Immun, 2020. 89(1).

38. Gopal, R., et al., S100A8/A9 proteins mediate neutrophilic inflammation and lung pathology during tuberculosis. Am J Respir Crit Care Med, 2013. 188(9): p. 1137–46.

39. Koyuncu, D., et al., CXCL1: A new diagnostic biomarker for human tuberculosis discovered using Diversity Outbred mice. PLoS Pathog, 2021. 17(8): p. e1009773.

40. Qiu, J., et al., ZBTB20-mediated titanium particle-induced peri-implant osteolysis by promoting macrophage inflammatory responses. Biomater Sci, 2020. 8(11): p. 3147–3163.

41. Pasqualini, R., et al., A study of the structure, function and distribution of beta 5 integrins using novel anti-beta 5 monoclonal antibodies. J Cell Sci, 1993. 105 **(Pt** **1**): p. 101–11.

42. Vogel, W., Discoidin domain receptors: structural relations and functional implications. The FASEB journal, 1999. 13(9001): p. S77–S82.

43. Mattiotti, A., et al., Follistatin-like 1 in development and human diseases. Cell Mol Life Sci, 2018. 75(13): p. 2339–2354.

44. Kuroda, Y., et al., A novel gene (FAM20B encoding glycosaminoglycan xylosylkinase) for neonatal short limb dysplasia resembling Desbuquois dysplasia. Clin Genet, 2019. 95(6): p. 713–717.

45. Koike, T., et al., FAM20B is a kinase that phosphorylates xylose in the glycosaminoglycan-protein linkage region. Biochem J, 2009. 421(2): p. 157–62.

46. Zhu, C., et al., Regulation of the Development and Function of B Cells by ZBTB Transcription Factors. Front Immunol, 2018. 9: p. 580.

47. Sutherland, A.P., et al., Zinc finger protein Zbtb20 is essential for postnatal survival and glucose homeostasis. Molecular and cellular biology, 2009. 29(10): p. 2804–2815.

48. Lightman, S.M., A. Utley, and K.P. Lee, Survival of Long-Lived Plasma Cells (LLPC): Piecing Together the Puzzle. Front Immunol, 2019. 10: p. 965.

49. Sun, Y., et al., Zbtb20 Restrains CD8 T Cell Immunometabolism and Restricts Memory Differentiation and Antitumor Immunity. J Immunol, 2020. 205(10): p. 2649–2666.

50. Schilling, D., M.R. Pittelkow, and R. Kumar, IEX-1, an immediate early gene, increases the rate of apoptosis in keratinocytes. Oncogene, 2001. 20(55): p. 7992–7997.

51. Im, H.-J., M.R. Pittelkow, and R. Kumar, Divergent Regulation of the Growth-promoting GeneIEX- 1 by the p53 Tumor Suppressor and Sp1. Journal of Biological Chemistry, 2002. 277(17): p. 14612–14621.

52. Arlt, A. and H. Schäfer, Role of the immediate early response 3 (IER3) gene in cellular stress response, inflammation and tumorigenesis. Eur J Cell Biol, 2011. 90(6-7): p. 545–52.

53. Harrison, D.E., et al., Genetically diverse mice are novel and valuable models of age-associated susceptibility to Mycobacterium tuberculosis. Immun Ageing, 2014. 11(1): p. 24.

54. Cyktor, J.C., et al., IL-10 inhibits mature fibrotic granuloma formation during Mycobacterium tuberculosis infection. J Immunol, 2013. 190(6): p. 2778–90.

55. Vesosky, B., et al., CCL5 participates in early protection against Mycobacterium tuberculosis. Journal of Leukocyte Biology, 2010. 87(6): p. 1153–65.

56. Ullman-Cullere, M.H. and C.J. Foltz, Body condition scoring: a rapid and accurate method for assessing health status in mice. Lab Anim Sci, 1999. 49(3): p. 319–23.

57. Tavolara, T.E., et al., Automatic discovery of clinically interpretable imaging biomarkers for Mycobacterium tuberculosis supersusceptibility using deep learning. EBioMedicine, 2020. 62: p. 103094.

58. Beamer, G.L., et al., Peripheral blood gamma interferon release assays predict lung responses and Mycobacterium tuberculosis disease outcome in mice. Clinical and Vaccine Immunology, 2008. 15(3): p. 474–83.

59. Keane, T.M., et al., Mouse genomic variation and its effect on phenotypes and gene regulation. Nature, 2011. 477(7364): p. 289–94.

60. Tavolara, T.E., et al., Deep learning predicts gene expression as an intermediate data modality to identify susceptibility patterns in Mycobacterium tuberculosis infected Diversity Outbred mice. EBioMedicine, 2021. 67: p. 103388.

61. Morgan, A.P., et al., The Mouse Universal Genotyping Array: From Substrains to Subspecies. G3 (Bethesda), 2015. 6(2): p. 263–79.

62. Broman, K.W., et al., R/qtl2: Software for Mapping Quantitative Trait Loci with High-Dimensional Data and Multiparent Populations. Genetics, 2019. 211(2): p. 495–502.

63. Cheng, R., et al., QTLRel: an R package for genome-wide association studies in which relatedness is a concern. BMC Genet, 2011. 12: p. 66.

64. Van Ooijen, J.W., LOD significance thresholds for QTL analysis in experimental populations of diploid species. Heredity (Edinb), 1999. 83 **(Pt** **5**): p. 613–24.

65. David Meyer, E.D., Kurt Hornik, Andreas Weingessel and Friedrich Leisch, e1071: Misc Functions of the Department of Statistics, Probability Theory Group (Formerly: E1071), TU Wien. 2021. p. R package.

66. Blake, J.A., et al., Mouse Genome Database (MGD): Knowledgebase for mouse-human comparative biology. Nucleic Acids Res, 2021. 49(D1): p. D981–D987.

67. Ahmed, M., et al., Immune correlates of tuberculosis disease and risk translate across species. Sci Transl Med, 2020. 12(528).

68. Kurtz, S.L., et al., The Diversity Outbred Mouse Population Is an Improved Animal Model of Vaccination against Tuberculosis That Reflects Heterogeneity of Protection. mSphere, 2020. 5(2).

69. Kus, P., M.N. Gurcan, and G. Beamer, Automatic Detection of Granuloma Necrosis in Pulmonary Tuberculosis Using a Two-Phase Algorithm: 2D-TB. Microorganisms, 2019. 7(12).

70. Niazi, M.K.K., G. Beamer, and M.N. Gurcan, An application of transfer learning to neutrophil cluster detection for tuberculosis: efficient implementation with nonmetric multidimensional scaling and sampling. SPIE Medical Imaging. Vol. 10581. 2018: SPIE.

71. Niazi, M.K.K., G. Beamer, and M.N. Gurcan, A computational framework to detect normal and tuberculosis infected lung from H and E-stained whole slide images. SPIE Medical Imaging. Vol. 10140. 2017: SPIE.

72. Sanchez, F., et al., Multigenic control of disease severity after virulent Mycobacterium tuberculosis infection in mice. Infect Immun, 2003. 71(1): p. 126–31.

73. Yan, B.S., et al., Genetic architecture of tuberculosis resistance in a mouse model of infection. Genes and Immunity, 2006. 7(3): p. 201–10.

74. Dodd, C.E. and L.S. Schlesinger, New concepts in understanding latent tuberculosis. Curr Opin Infect Dis, 2017. 30(3): p. 316–321.

75. Behr, M.A., P.H. Edelstein, and L. Ramakrishnan, Revisiting the timetable of tuberculosis. Bmj, 2018. 362: p. k2738.

76. Loddenkemper, R., M. Lipman, and A. Zumla, Clinical Aspects of Adult Tuberculosis. Cold Spring Harb Perspect Med, 2015. 6(1): p. a017848.

77. Hunter, R.L., et al., Pathogenesis of post primary tuberculosis: immunity and hypersensitivity in the development of cavities. Ann Clin Lab Sci, 2014. 44(4): p. 365–87.

78. Naranbhai, V., The Role of Host Genetics (and Genomics) in Tuberculosis. Microbiol Spectr, 2016. 4(5).

79. Abel, L., et al., Human genetics of tuberculosis: a long and winding road. Philos Trans R Soc Lond B Biol Sci, 2014. 369(1645): p. 20130428.

80. Kondratieva, E., et al., Host genetics in granuloma formation: human-like lung pathology in mice with reciprocal genetic susceptibility to M. tuberculosis and M. avium. PLoS One, 2010. 5(5): p. e10515.

81. Bustamante, J., et al., Mendelian susceptibility to mycobacterial disease: genetic, immunological, and clinical features of inborn errors of IFN-gamma immunity. Semin Immunol, 2014. 26(6): p. 454–70.

82. Qu, H.Q., S.P. Fisher-Hoch, and J.B. McCormick, Molecular immunity to mycobacteria: knowledge from the mutation and phenotype spectrum analysis of Mendelian susceptibility to mycobacterial diseases. Int J Infect Dis, 2011. 15(5): p. e305–13.

83. Cottle, L.E., Mendelian susceptibility to mycobacterial disease. Clin Genet, 2011. 79(1): p. 17–22.

84. Kramnik, I., Genetic dissection of host resistance to Mycobacterium tuberculosis: the sst1 locus and the Ipr1 gene. Curr Top Microbiol Immunol, 2008. 321: p. 123–48.

85. Tosh, K., et al., Variants in the SP110 gene are associated with genetic susceptibility to tuberculosis in West Africa. Proc Natl Acad Sci U S A, 2006. 103(27): p. 10364–10368.

86. Pan, H., et al., Ipr1 gene mediates innate immunity to tuberculosis. Nature, 2005. 434(7034): p. 767–72.

87. Kramnik, I., P. Demant, and B.B. Bloom, Susceptibility to tuberculosis as a complex genetic trait: analysis using recombinant congenic strains of mice. Novartis Foundation symposium, 1998. 217: p. 120–31; discussion 132-7.

88. Apt, A.S., et al., Distinct H-2 complex control of mortality, and immune responses to tuberculosis infection in virgin and BCG-vaccinated mice. Clinical and experimental immunology, 1993. 94(2): p. 322–9.

89. Barrera, L.F., et al., I-A beta gene expression regulation in macrophages derived from mice susceptible or resistant to infection with M. bovis BCG. Mol Immunol, 1997. 34(4): p. 343–55.

90. Logunova, N., et al., The QTL within the H2 Complex Involved in the Control of Tuberculosis Infection in Mice Is the Classical Class II H2-Ab1 Gene. PLoS Genet, 2015. 11(11): p. e1005672.

91. Scott, N.R., et al., S100A8/A9 regulates CD11b expression and neutrophil recruitment during chronic tuberculosis. J Clin Invest, 2020. 130(6): p. 3098–3112.

92. Yang, H., et al., On the subspecific origin of the laboratory mouse. Nat Genet, 2007. 39(9): p. 1100–7.

93. Jiao, L., et al., A Novel Genetic Variation in NCF2, the Core Component of NADPH Oxidase, Contributes to the Susceptibility of Tuberculosis in Western Chinese Han Population. DNA Cell Biol, 2020. 39(1): p. 57–62.

94. Cooper, A.M., et al., Transient loss of resistance to pulmonary tuberculosis in p47(phox-/-) mice. Infect Immun, 2000. 68(3): p. 1231–4.

95. Wu, J., et al., FAM20B-catalyzed glycosaminoglycans control murine tooth number by restricting FGFR2b signaling. BMC Biol, 2020. 18(1): p. 87.

96. Saiyin, W., et al., Inactivation of FAM20B causes cell fate changes in annulus fibrosus of mouse intervertebral disc and disc defects via the alterations of TGF-beta and MAPK signaling pathways. Biochim Biophys Acta Mol Basis Dis, 2019. 1865(12): p. 165555.

97. Ishikawa, H.O., et al., Four-jointed is a Golgi kinase that phosphorylates a subset of cadherin domains. Science, 2008. 321(5887): p. 401–404.

98. Perez, T.D. and W.J. Nelson, Cadherin adhesion: mechanisms and molecular interactions. Handb Exp Pharmacol, 2004(165): p. 3–21.

99. Zygiel, E.M. and E.M. Nolan, Transition Metal Sequestration by the Host-Defense Protein Calprotectin. Annu Rev Biochem, 2018. 87: p. 621–643.

100. Wang, S., et al., S100A8/A9 in Inflammation. Front Immunol, 2018. 9: p. 1298.

101. Dale, C.S., et al., Analgesic properties of S100A9 C-terminal domain: a mechanism dependent on calcium channel inhibition. Fundam Clin Pharmacol, 2009. 23(4): p. 427–38.

102. Dale, C.S., et al., Effect of the C-terminus of murine S100A9 protein on experimental nociception. Peptides, 2006. 27(11): p. 2794–802.

103. Hudson, B.I. and M.E. Lippman, Targeting RAGE Signaling in Inflammatory Disease. Annu Rev Med, 2018. 69: p. 349–364.

104. Garcia, V., Y.R. Perera, and W.J. Chazin, A Structural Perspective on Calprotectin as a Ligand of Receptors Mediating Inflammation and Potential Drug Target. Biomolecules, 2022. 12(4).

105. Ryckman, C., et al., Proinflammatory activities of S100: proteins S100A8, S100A9, and S100A8/A9 induce neutrophil chemotaxis and adhesion. The Journal of Immunology, 2003. 170(6): p. 3233–3242.

106. Gopal, R., et al., S100A8/A9 proteins mediate neutrophilic inflammation and lung pathology during tuberculosis. American journal of respiratory and critical care medicine, 2013. 188(9): p. 1137–1146.

107. Liu, Q., et al., High levels of plasma S100A9 at admission indicate an increased risk of death in severe tuberculosis patients. Journal of Clinical Tuberculosis and Other Mycobacterial Diseases, 2021. 25: p. 100270.

108. Li, H.M., et al., Altered NCF2, NOX2 mRNA Expression Levels in Peripheral Blood Mononuclear Cells of Pulmonary Tuberculosis Patients. Int J Gen Med, 2021. 14: p. 9203–9209.

109. Leung, K., *(99m)Tc-Diamine dioxime-Lys-Cys-Arg-Gly-Asp-Cyc-Phe-Cys-polyethylene glycol*, in *Molecular Imaging and Contrast Agent Database (MICAD)*. 2004, National Center for Biotechnology Information (US): Bethesda (MD).

110. Lin, Z., et al., Integrin-β5, a miR-185-targeted gene, promotes hepatocellular carcinoma tumorigenesis by regulating β-catenin stability. Journal of Experimental & Clinical Cancer Research, 2018. 37: p. 1–13.

111. Shi, W., et al., Integrin beta5 enhances the malignancy of human colorectal cancer by increasing the TGF-beta signaling. Anticancer Drugs, 2021. 32(7): p. 717–726.

112. Zhuang, H., et al., Characterization of the prognostic and oncologic values of ITGB superfamily members in pancreatic cancer. J Cell Mol Med, 2020. 24(22): p. 13481–13493.

113. Nurzat, Y., et al., Identification of Therapeutic Targets and Prognostic Biomarkers Among Integrin Subunits in the Skin Cutaneous Melanoma Microenvironment. Front Oncol, 2021. 11: p. 751875.

114. Zhang, L.Y., et al., Integrin Beta 5 Is a Prognostic Biomarker and Potential Therapeutic Target in Glioblastoma. Front Oncol, 2019. 9: p. 904.

115. Ouchi, N., et al., DIP2A functions as a FSTL1 receptor. J Biol Chem, 2010. 285(10): p. 7127–34.

116. Jin, X., et al., Fstl1 promotes glioma growth through the BMP4/Smad1/5/8 signaling pathway. Cellular Physiology and Biochemistry, 2017. 44(4): p. 1616–1628.

117. Murakami, K., et al., Follistatin-related protein/follistatin-like 1 evokes an innate immune response via CD14 and toll-like receptor 4. FEBS letters, 2012. 586(4): p. 319–324.

118. Horak, M., et al., Follistatin-like 1 and its paralogs in heart development and cardiovascular disease. Heart Fail Rev, 2022. 27(6): p. 2251–2265.

119. Zhang, Y., et al., Circ-WDR27 regulates mycobacterial vitality and secretion of inflammatory cytokines in Mycobacterium tuberculosis-infected macrophages via the miR-370-3p/FSTL1 signal network. J Biosci, 2022. 47.

120. Zhang, W., et al., Identification and characterization of DPZF, a novel human BTB/POZ zinc finger protein sharing homology to BCL-6. Biochem Biophys Res Commun, 2001. 282(4): p. 1067–73.

121. Xie, Z., et al., Zinc finger protein ZBTB20 is a key repressor of alpha-fetoprotein gene transcription in liver. Proceedings of the National Academy of Sciences, 2008. 105(31): p. 10859–10864.

122. Xie, Z., et al., Zbtb20 is essential for the specification of CA1 field identity in the developing hippocampus. Proceedings of the National Academy of Sciences, 2010. 107(14): p. 6510–6515.

123. Zhang, Y., et al., *The zinc finger protein ZBTB20 regulates transcription of fructose-1*, *6- bisphosphatase 1 and β cell function in mice*. Gastroenterology, 2012. 142(7): p. 1571–1580.e6.

124. Liu, X., et al., Zinc finger protein ZBTB20 promotes Toll-like receptor-triggered innate immune responses by repressing IκBα gene transcription. Proc Natl Acad Sci U S A, 2013. 110(27): p. 11097–102.

125. Arora, V., et al., *Primrose Syndrome*, in *GeneReviews*, M.P. Adam, et al., Editors. 1993, University of Washington, Seattle: Seattle (WA).

126. Kim, S.H., et al., Discoidin domain receptor 1 mediates collagen-induced nitric oxide production in J774A.1 murine macrophages. Free Radic Biol Med, 2007. 42(3): p. 343–52.

127. Wang, H., et al., DDR1 activation in macrophage promotes IPF by regulating NLRP3 inflammasome and macrophage reaction. Int Immunopharmacol, 2022. 113(Pt A): p. 109294.

128. Yeh, Y.C., H.H. Lin, and M.J. Tang, Dichotomy of the function of DDR1 in cells and disease progression. Biochim Biophys Acta Mol Cell Res, 2019. 1866(11): p. 118473.

129. Taciak, B., et al., Evaluation of phenotypic and functional stability of RAW 264.7 cell line through serial passages. PLoS One, 2018. 13(6): p. e0198943.

130. Ralph, P., J. Prichard, and M. Cohn, Reticulum cell sarcoma: an effector cell in antibody-dependent cell-mediated immunity. J Immunol, 1975. 114(2 pt 2): p. 898–905.

131. Zhang, X., et al., DDR1 promotes hepatocellular carcinoma metastasis through recruiting PSD4 to ARF6. Oncogene, 2022. 41(12): p. 1821–1834.

132. Su, H., et al., Collagenolysis-dependent DDR1 signalling dictates pancreatic cancer outcome. Nature, 2022. 610(7931): p. 366–372.

133. Ma, R., et al., Discoidin domain receptors (DDRs): Potential implications in periodontitis. J Cell Physiol, 2022. 237(1): p. 189–198.

134. Moll, S., et al., DDR1 role in fibrosis and its pharmacological targeting. Biochim Biophys Acta Mol Cell Res, 2019. 1866(11): p. 118474.

135. Sirvent, A., et al., New functions of DDR1 collagen receptor in tumor dormancy, immune exclusion and therapeutic resistance. Front Oncol, 2022. 12: p. 956926.

136. Marchioro, H.Z., et al., Update on the pathogenesis of vitiligo. An Bras Dermatol, 2022. 97(4): p. 478–490.

137. Vilella, E., et al., Expression of DDR1 in the CNS and in myelinating oligodendrocytes. Biochim Biophys Acta Mol Cell Res, 2019. 1866(11): p. 118483.

138. Wu, M., Roles of the stress-induced gene IEX-1 in regulation of cell death and oncogenesis. Apoptosis, 2003. 8: p. 11–18.

139. Arlt, A. and H. Schäfer, Role of the immediate early response 3 (IER3) gene in cellular stress response, inflammation and tumorigenesis. European journal of cell biology, 2011. 90(6-7): p. 545–552.

140. Arlt, A., et al., Expression of the NF-κB target gene IEX-1 (p22/PRG1) does not prevent cell death but instead triggers apoptosis in HeLa cells. Oncogene, 2001. 20(1): p. 69–76.

141. Sebens Müerköster, S., et al., The apoptosis-inducing effect of gastrin on colorectal cancer cells relates to an increased IEX-1 expression mediating NF-κB inhibition. Oncogene, 2008. 27(8): p. 1122–1134.

142. Akilov, O.E., et al., Enhanced susceptibility to Leishmania infection in resistant mice in the absence of immediate early response gene X-1. J Immunol, 2009. 183(12): p. 7994–8003.

143. Leisching, G., et al., RNAseq reveals hypervirulence-specific host responses to M. tuberculosis infection. Virulence, 2017. 8(6): p. 848–858.

144. Coultas, L., K. Chawengsaksophak, and J. Rossant, Endothelial cells and VEGF in vascular development. Nature, 2005. 438(7070): p. 937–45.

145. Ferrara, N. and H.P. Gerber, The role of vascular endothelial growth factor in angiogenesis. Acta Haematol, 2001. 106(4): p. 148–56.

146. Ferrara, N. and K. Alitalo, Clinical applications of angiogenic growth factors and their inhibitors. Nat Med, 1999. 5(12): p. 1359–64.

147. Shibuya, M., Vascular Endothelial Growth Factor (VEGF) and Its Receptor (VEGFR) Signaling in Angiogenesis: A Crucial Target for Anti- and Pro-Angiogenic Therapies. Genes Cancer, 2011. 2(12): p. 1097–105.

148. Davuluri, K.S., et al., Stimulated expression of ELR+ chemokines, VEGFA and TNF-AIP3 promote mycobacterial dissemination in extrapulmonary tuberculosis patients and Cavia porcellus model of tuberculosis. Tuberculosis (Edinb), 2022. 135: p. 102224.

149. Delemarre, E.M., et al., Serum Biomarker Profile Including CCL1, CXCL10, VEGF, and Adenosine Deaminase Activity Distinguishes Active From Remotely Acquired Latent Tuberculosis. Front Immunol, 2021. 12: p. 725447.

150. Golubinskaya, E.P., et al., Dysregulation of VEGF-dependent angiogenesis in cavernous lung tuberculosis. Pathophysiology, 2019. 26(3-4): p. 381–387.

151. Harding, J.S., et al., VEGF-A from Granuloma Macrophages Regulates Granulomatous Inflammation by a Non-angiogenic Pathway during Mycobacterial Infection. Cell Rep, 2019. 27(7): p. 2119–2131.e6.

152. Tao, R.-H., et al., Opposite effects of alternative TZF spliced variants on androgen receptor. Biochemical and biophysical research communications, 2006. 341(2): p. 515–521.

153. Pioli, P.D., et al., Zfp318 regulates IgD expression by abrogating transcription termination within the Ighm/Ighd locus. J Immunol, 2014. 193(5): p. 2546–53.

154. Enders, A., et al., Zinc-finger protein ZFP318 is essential for expression of IgD, the alternatively spliced Igh product made by mature B lymphocytes. Proceedings of the National Academy of Sciences, 2014. 111(12): p. 4513–4518.

155. Schwickert, T.A., et al., Ikaros prevents autoimmunity by controlling anergy and Toll-like receptor signaling in B cells. Nat Immunol, 2019. 20(11): p. 1517–1529.

